# Maternal obesogenic diet disrupts mid-gestation decidual immune and vascular homeostasis without impairing spiral artery remodelling

**DOI:** 10.1101/2024.09.10.612284

**Authors:** Christian J. Bellissimo, Erica Yeo, Tatiane A. Ribeiro, Patrycja A. Jazwiec, Chethana Ellewela, Jaskiran Bains, Ali A. Ashkar, Alexander G. Beristain, Dawn M.E. Bowdish, Deborah M. Sloboda

## Abstract

Excess maternal adiposity (i.e., overweight and obesity) during pregnancy has been linked to impaired uteroplacental perfusion, compromised placental development, and a higher risk of adverse pregnancy outcomes. Owing to the nature of chronic inflammation and immune dysregulation accompanying excess adiposity, disruption of leukocyte-mediated tissue remodelling and immunoregulation within the decidua have emerged as likely drivers contributing to suboptimal placental function in pregnancies impacted by maternal overweight or obesity. However, the impacts of excess adiposity on major populations of innate lymphoid cells (ILCs) and macrophages which orchestrate these processes and the environment that these cells occupy remain vastly understudied. Here, we used a mouse model of chronic high-fat, high-sucrose (HFHS) diet-feeding to characterize the impacts of an obesogenic milieu on decidual immune dynamics during placental development at mid-gestation (E10.5). HFHS pregnancies exhibited marked increases in total decidual leukocyte abundance, driven by population-level increases in tissue-resident and conventional NK cells, and MHC-II^+^ macrophages. This was not associated with abnormalities in implantation site morphology or decidual spiral artery remodelling but was coincident with histological patterns of local inflammation. In line with this, expression of canonical proinflammatory cytokines and chemokines were moderately upregulated in bulk decidual tissue of HFHS dams. This was accompanied by more potent elevations in multiple mediators of angiogenesis, endothelial activation, and coagulation in HFHS decidual tissue. Collectively, these findings point towards pathological vascular inflammation and possibly dysregulated decidual angiogenesis in the first half of pregnancy as factors predisposing to reduced placental efficiency, malperfusion, and inflammation seen in pregnancies affected by maternal overweight and obesity.

## INTRODUCTION

The placenta is a critical maternal-fetal interface throughout pregnancy, facilitating nutrient and gas exchange, manipulating maternal physiologic adaptations to pregnancy, and forming a protective barrier against xenobiotics, pathogens, and maternal allorecognition (1). Normal placental development is, in part, dependent upon a carefully regulated balance of canonically proinflammatory and immunoregulatory signals derived from resident immune cell populations in the uterine mucosa (decidua) (2, 3). These leukocytes orchestrate both structural remodelling of the decidua and maintenance of a local tolerogenic environment, ensuring optimal placental function and fetal growth.

Decidual leukocyte populations are largely made up of innate lymphoid cells (ILCs) and myeloid cells (∼80 – 90%) in the first half of pregnancy, with decidual NK (dNK) cell and macrophages predominating these populations, respectively (4–6). Decidual macrophages and dNK cells cooperate in the remodelling of decidual spiral arteries, producing cytokines and matrix remodelling enzymes that trigger vascular media destruction, as well as chemotactic factors which regulate the differentiation of invasive trophoblast cell lineages and their interstitial and endovascular migration (4, 7–10). Growth factors produced by decidual leukocytes—particularly dNK cells—also stimulate decidual angiogenesis and placental growth (11, 12).

In addition to their trophic functions, decidual leukocytes and stromal cells are a source of immunomodulatory cytokines that counterbalance inflammatory byproducts of tissue remodelling and create a local bias towards regulatory immunity. As the primary antigen-presenting cells within the decidua, macrophages are implicated in regulatory T (T_reg_) differentiation, suppressing maternal allorecognition of antigenically foreign trophoblast by the adaptive immune system (13, 14). Despite their tolerogenic bias, most uterine-resident leukocytes retain the capacity to induce cytotoxic responses under inflammatory conditions (15–17). While important for host defence, excessive production of inflammatory effectors can have pathological impacts on placental development and function, leading to complications up to and including pregnancy loss (18–20). Conversely, impaired recruitment, differentiation, or activation of decidual immune cell populations can disrupt the progression of decidual vascular transformation, resulting in uteroplacental malperfusion, reperfusion injury, and activation of inflammatory and hypoxic signalling cascades (21–25). These changes can lead to placental insufficiency and underlie many obstetrical disorders preterm birth, fetal growth restriction, stillbirth, and preeclampsia (22, 26, 27). Even in the absence of overt clinical complications, suboptimal placental function has been linked to postnatal programming of offspring susceptibility to multiple chronic diseases (1, 28). As such, pre-existing health conditions associated with excessive maternal inflammation or immune dysregulation are linked to an increased risk of adverse pregnancy outcomes, perinatal morbidity, and poorer maternal and child health postnatally.

Maternal excess adiposity is the most common chronic condition affecting pregnant people in North America, with nearly half of all individuals entering gestation overweight (BMI 25.0 – 29.9) or obese (BMI ≥30) (29, 30). The risk of most obstetrical disorders increases alongside maternal BMI, including an increased frequency of pregnancy loss, preterm delivery, stillbirth, and preeclampsia (31, 32). Extremes in fetal growth are also common, with more term infants being born macrosomic or, in the case of severely obese individuals, small for gestational age (31–36). In all cases, excess maternal adiposity has been linked to reduced placental efficiency, impaired fetoplacental vascular development, and an increased burden of inflammatory and malperfusion-related pathologies (37–40). Systemic inflammation and impaired effector functions of immune cells are thought to play a prominent role in obesity-associated placental dysfunction, potentiating adverse perinatal and long-term health outcomes (41–43). However, despite several studies documenting peripheral immune and inflammatory changes in both pregnant women and animal models with obesity, (44–49) only a few recent studies have directly profiled the impacts of excess adiposity on the uterine immune environment during placental development (50–54).

These studies have reported reduced frequencies and dysregulated activity and cytokine production of first-trimester dNK cells in pregnancies complicated by maternal obesity (51, 52). These changes were also accompanied by deficient early-phase remodelling of spiral arteries, potentially contributing to obesity-associated placental malperfusion (51). Similar remodelling impairments have been reported in animal models of diet-induced excess adiposity (53, 55). However, decidual immunophenotypic assessments in these studies were either absent or limited to the characterization of dNK cells, with some studies in rodents exhibiting discrepancies with human findings (51–53), or lacking a broader survey of the decidual immune and angiogenic environment (54, 55). Like other metabolic tissues (56–60), increased densities of uterine macrophages have been reported both before conception and at term in humans and rodents with obesity (17, 61, 62). However, phenotypic characterization of these cells, their microenvironment, and associated uteroplacental outcomes are largely unclear.

In this study, we used an established model of high-fat, high-sucrose (HFHS) feeding in mice (53, 63, 64) to profile the combined impacts of excess adiposity and a maternal obesogenic milieu on the uterine immune environment during the peak of decidual vascular remodelling. We hypothesized that HFHS diet-induced excess adiposity would impair the progression of spiral artery remodelling, owing to reduced numbers of dNK cells and inflammatory macrophage skewing in the mid-gestation decidua. Here, we report that while diet-induced excess adiposity does not impact early-phase vascular media destruction or angiogenesis in spiral arteries, the numbers of both decidual macrophages and dNK are increased at mid-gestation in HFHS dams. This was accompanied by modest elevations in pro-inflammatory mediators and pronounced dysregulation of factors involved in angiogenesis and coagulation. Overall, these findings reveal distinct cellular and molecular shifts during early placental development that may represent drivers of and/or responses to uteroplacental malperfusion, local inflammation, and placental dysfunction seen in pregnancies affected by maternal excess adiposity.

## METHODS

### Sex as a biological variable

This work aimed to disentangle the impacts of excess adiposity in the maternal lineage on the uterine immune environment in early pregnancy. It is now clear that metabolic dysfunction in both parents uniquely contributes to alterations in uterine immune cell composition and function in the periconceptional period (17, 51, 65). Thus, it is unclear whether the findings we report are representative of the scenario in which both parents are overweight or obese. Where possible, we have stratified histologic and molecular analyses of uteroplacental tissues by conceptus sex to determine whether the outcomes of maternal metabolic dysfunction on vascular remodelling and the uterine immune environment are sex-dependent.

### Animal model and sample collection

Virgin C57BL/6J female mice were purchased from Jackson Laboratories at 7 weeks of age and acclimated for one week in the McMaster Central Animal Facility before experimental use. Non-pregnant females were housed in groups of four under specific-pathogen-free conditions in individually vented cages with controlled temperature and humidity (22 – 24°C, 40 – 60% RH) on a 12-hour light/dark cycle. Food and water were provided *ad libitum*. At baseline (8 weeks of age), and 10 weeks following diet onset, female mice were fasted for six hours and subject to oral glucose tolerance tests to assess glycemic control (detailed below). Following the initial OGTT, females were randomized to receive either an obesogenic high-fat, high-sucrose diet (HFHS, 4.73 kCal/g: 45% fat, 17% sucrose, 18% carbohydrate, 20% protein; Research Diets D12451) or maintained on a fixed-formula rodent chow diet (CON, 3.0 kCal/g: 17% fat, 54% carbohydrate, 29% protein; Teklad 22/5 Rodent Diet, Envigo cat. 8640) as a control group, ensuring baseline body weight, fasting glucose or oral glucose tolerance was similar between groups. A comparison of dietary macro- and micronutrients is outlined in Supplementary Table 1. Following 12 weeks of dietary intervention, CON and HFHS females were time-mated with young (14- to 20-weeks old) C57BL/6J studs of proven fertility overnight (19:00 – 08:00 hrs). Successful mating was identified by the presence of a copulatory plug in the morning following pairing and designated E0.5. Pregnant dams were singly housed and allowed to carry pregnancies to E10.5. On E10.5, dams were fasted for six hours (03:00-0:900h), and blood was collected for blood glucose measurement and serum isolation (detailed below). Immediately following this, dams were euthanized by cervical dislocation for the collection of reproductive and metabolic tissues. As not all pairings resulted in copulation, and not all successful mating generated a pregnancy, the length of preconception dietary intervention varied. Females from both groups who did not conceive by 17 weeks of diet intervention were excluded from this study to limit variance in outcomes due to prolonged diet exposure (46, 60).

Uterine tissues were immediately isolated following laparotomy, and the number and position of implantation sites counted. The presence of embryonic resorptions or markedly pale/underdeveloped implantation sites was recorded and excluded from downstream analyses. A ligated segment of four implantation sites was excised from the uterus and fixed by immersion in 4% paraformaldehyde in phosphate-buffered saline pH 7.4 for 24 hours at 4°C and processed for paraffin embedding and histology, as previously described (66). The remaining implantation sites were individually segmented through inter-implantation regions, and three to four implantation sites per dam were reserved for decidual immunophenotyping. Where more than 7 – 8 viable implantation sites in a given pregnancy, remaining implantations were tissue microdissected to isolate myometrial, decidual, and placental tissue (67, 68). The resulting tissue layers and embryonic tissue were separately snap-frozen in liquid nitrogen and stored at -80°C until further processing.

### Measures of glycemic control

Glycemic control in non-pregnant females was assessed by an oral glucose challenge following a six-hour fast (03:00 – 09:00 hrs). Mice were administered a bolus of 30% w/v solution of D-glucose in isotonic saline at a dose of 2g/kg of body weight by oral gavage. Blood was sampled using a controlled tail vein bleed. Blood glucose was recorded at 0-, 15-, 30-, 60-, 90-, and 120-minutes after gavage using a handheld glucometer (Accu-Check Aviva, Roche Diagnostics). Glucose tolerance was measured by calculating the incremental area under the glucose curve (iAUC), using the fasting blood glucose of individual mice as a baseline from which glucose clearance from circulation was measured over 120 minutes (69).

Thirty minutes before gavage, 150 μl of whole blood was collected in heparinized micro-hematocrit tubes (Fisher Scientific) for measurement of serum insulin levels. Blood was allowed to clot for 30 minutes at room temperature before isolation of serum by centrifugation of samples at 1,000 × g for 10 minutes. Serum insulin concentrations were determined using a commercial ultra-sensitive ELISA (Toronto Biosciences cat. #32380) according to manufacturer’s instructions. Samples with absorbance at background level were set to half of the lower limit of detection (0.0125 ng/ml).

### Decidual immunophenotyping

Single cell digests of decidual tissue were generated as previously described (53, 68, 70, 71), with some modification. In brief, mesometrial decidual tissue with the placenta *in situ* was dissected from three to four implantation sites per pregnancy in ice-cold HBSS supplemented with 15 mM HEPES and 10% FCS (Wisent Biosciences). Isolated decidual tissues were then rinsed with RPMI-1640 basal media, pooled, and minced until tissue fragments were <1mm^2^ in size. Minced tissue was transferred to a final volume of 2.5 mL of enzymatic tissue digestion media (50 μg/mL Liberase TM (0.28 Wünsch units; Roche Diagnostics) and 50 μg/ml Dnase I in RPMI). Tissues were digested at 37°C with gentle trituration every 10 minutes, for 30 minutes total. Suspensions were filtered through a 70-μm nylon cell sieve into 1mL of FBS to quench the enzymatic reaction. Small pieces of remaining tissue were macerated with a syringe plunger over the filter mesh followed by washing several times with RPMI to disaggregate remaining cells. Suspensions were topped up to a final volume of 20 mL with RPMI (5% v/v FBS) and pelleted by centrifugation at 350×g for 10 minutes. Cell pellets were resuspended in ACK lysis buffer (ThermoFisher Scientific) for 90 seconds at room temperature to deplete red blood cells. The suspension was diluted 1:10 with ice-cold phosphate-buffered saline (PBS, Wisent Biosciences), pelleted, washed, resuspended in 250 μl of PBS and immediately subject to counting and staining for flow cytometry analysis.

Separation of decidual and myometrial tissue, including the mesometrial lymphoid aggregate of pregnancy (MLAp), was confirmed by the presence of both NK cells and macrophages in paired flow cytometry analyses in each sample and a distinctive staining profile (Supplementary Figure 1). As cells of the MLAp do not associate with actively transforming spiral arteries and the efficacy of tissue digestion varied widely for myometrial samples, they were not included for analysis in this study.

Total suspension volume (resuspension PBS + residual liquid) was recorded for later absolute count calculations. Cells were counted using a hemocytometer and suspension volumes containing ∼5×10^5^ cells/pregnancy were taken for cell staining in a 96-well plate, and the remaining cells were equally divided for isotype or flow-minus-one (FMO) staining controls. Cells were pelleted and resuspended in PBS with anti-CD16/32 diluted 1:25 (Fc block clone 2.4G2; BD Biosciences) for 15 minutes at 4°C to block non-specific staining. Following this, an equal volume of 2× concentrated antibody staining cocktail in PBS was added directly to the cell suspension (Supplementary Table 2). Cells were incubated for 30 minutes at 4°C, protected from light. Following surface staining, cell suspensions were pelleted, washed, and underwent fixation, permeabilization, and intranuclear staining for Eomesodermin using the FOXP3/transcription factor staining kit (eBioscience) following manufacturer’s instructions. Stained samples were then washed and resuspended in equal volumes of PBS. CountBright Absolute Counting Beads (Invitrogen) were added to each sample and filtered through a 70-μm cell sieve immediately before acquisition on a Beckman Coulter CytoFlex LX, using single stained VersaComp antibody capture beads to generate compensation matrices. Data were analysed in FlowJo v10.7.1 (BD), using isotype and FMO controls to define gating cut-offs. Absolute counts of macrophage and dILC populations were calibrated across multiple sample preps by multiplying the raw event counts of each population by a sample-specific calibration factor. This product was then normalized to the total amount of tissue used for digestion (both total tissue weight and the total number of digested implantation sites). Calibration factors were calculated as follows:

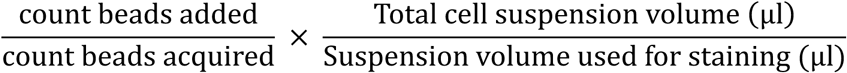

### Genotyping for sex determination

Sex determination of conceptuses occupying frozen or fixed implantation sites was performed by duplex PCR-genotyping of embryonic tissues for the Y-linked Sry (male sex) and autosomal Fabp2 gene loci (internal amplification control) using genotyping primers designed by the Mouse Genetics Core at Washing University School of Medicine at St. Louis (https://mousegeneticscore.wustl.edu/items/pcr-genotyping-primer-pairs/, Supplementary Table 3). For implantation site sections from paraffin blocks, genomic DNA (gDNA) was extracted using the ReliaPrep FFPE gDNA miniprep system (Promega). For tissues used in molecular analyses, snap-frozen embryos isolated from the implantation sites were used for gDNA extraction with the DNA FastExtract Kit (Wisent Biosciences). Both were used according to manufacturer’s instructions. Amplicons were resolved by electrophoresis on a 1% agarose TAE gel with SYBRSafe (ThermoFisher Scientific) and imaged under UV light using a ChemiDoc MP Imaging System (1708280, BioRad). The presence of two amplicon bands at 273 bp (*Sry*) and 194 bp (*Fapb2*) indicated male sex, while a single band at 194 bp indicated female sex.

### Histology

Formaldehyde-fixed, paraffin-embedded tissues were subject to systematic serial sectioning using a rotary microtome at a thickness of 7-μm for downstream histological measures. Sections 250 to 400 (i.e., within the mid-sagittal plane) were collected for downstream histological analysis. Whole cross-section images of stained samples were acquired by slide scanning and image stitching at 20× magnification using a Nikon NiE Eclipse Microscope with NIS Elements software (v.5.20.02). All measurements of acquired images were performed by an observer blinded to the sample ID (CJB). Experiment-specific antigen retrieval conditions and antibodies for immunofluorescent studies are listed in Supplementary Table 4. Following antigen retrieval, samples were washed in PBS and blocked with 10% normal goat serum in PBS with 0.1% Tween20 (PBS-T) for 1 hour at room temperature. Sections were incubated with primary antibodies diluted in blocking buffer overnight at 4°C. The following day, samples were washed and incubated with fluorophore- or biotin-conjugated secondary detection reagents in PBS-T for 1 hour at room temperature. For immunofluorescence experiments with tertiary signal amplification (F4/80), sections were washed and incubated with fluorophore-conjugated Streptavidin diluted in PBS-T. Samples were counterstained with DAPI (1:5000, Molecular Probes) and mounted using Prolong Gold anti-fade mounting medium (Invitrogen). Positive staining was identified based on comparisons to sections stained with an isotype-matched non-specific primary antibody.

Morphometric assessments of spiral artery wall and lumen areas were performed on duplicate midline H&E-stained tissue sections, with each replicate 98-µm apart and containing a high density of arterial lumen. Wall and lumen areas were defined according to previously published criteria (53, 66, 68). In brief, spiral arteries were differentiated from venous vessels based on endothelial morphology (rounded appearance of the vascular endothelium and notable vessel wall (8, 68, 72, 73)). The five largest and roundest arterial vessels within each section were measured. Vessels that were greater than 1.5× wider along their major versus their minor axes were excluded as they were likely sagittal profiles, and where multiple arterial profiles were directly contiguous with one another, only one representative vessel was selected for analysis if more than five sufficiently large vessels meeting size and shape criteria were met, to minimize repeated measurement of different segments of the same spiral artery. Areas were measured using Nikon NIS Elements AR Analysis software.

Metrics of vascular media destruction and arterial angiogenesis were examined using triple immunofluorescent staining for α-SMA, Vimentin, and Ki-67 to identify vascular smooth muscle cells (VSMCs), stromal and endothelial cells, and actively proliferating cells, respectively. The endothelial perimeter was measured in all arterial profiles with a defined lumen present within the section by tracing the outer boundary of the endothelial layer (luminal VIM^+^ α-SMA^-^ cells). The progression of vascular remodelling was determined by dividing the cumulative length of the endothelial perimeter that was directly contoured by α-SMA staining. An estimate of spiral artery angiogenesis was made by measuring the number of DAPI-stained nuclei within the VIM^+^ endothelial layer that showed distinct staining of Ki-67. All measures were manually recorded using NIS Elements Analysis software (v.5.20.02).

### Decidual and serum protein analyses

Halves of frozen decidual tissue were homogenized using a bead mill with ceramic beads in 450 μl of extraction buffer (50 mM Tris, 150 mM NaCl, 20 mM NaF) supplemented with Pierce Protease Inhibitor Cocktail (ThermoFisher Scientific) containing Aprotinin, Bestatin, E-64, Leupeptin, AEBSF, Pepstatin A, and EDTA. Following homogenization, 50 μl of extraction buffer supplemented with 10% v/v Triton X-100 was added to each sample for a final concentration of 1% Triton X-100 in 500 μl. Crude lysates were vortexed, incubated on ice for 10 minutes, and clarified by centrifugation at 12,000×g for 15 minutes at 4°C to remove cellular debris and nuclei. Supernatants were collected, aliquoted, flash-frozen, and stored at -80°C until further use.

In both tissue lysates and pregnant maternal serum, a 12-plex panel of cytokines and growth factors with known roles in angiogenesis, inflammation and placental development were measured using the Milliplex Mouse Angiogenesis/Growth Factor Magnetic Bead Panel (MAGPMAG-24K, Millipore Sigma), and included the following analytes: leptin, EGF, IL-6, Endoglin (ENG), Endothelin-1, HGF, Placental growth factor (PlGF), CXCL1, CCL2, Prolactin, VEGF-A, and TNF Samples were thawed, clarified by centrifugation at 12,000×g for 15 minutes and run according to the manufacturer’s instructions. For decidual lysates, individual clarified samples were diluted 1:2 in assay buffer; for standard curves and assay controls, the serum matrix was replaced with complete lysate extraction buffer. Data were acquired using a Luminex MAGPIX-xMAP system. Following data acquisition, each sample was inspected for quality control. Any analyte measure for a given sample with fewer than 30 beads acquired was not considered reliable and was excluded from the analysis. In tissue samples, the raw concentration of each analyte detected was normalized to the amount of total protein in each sample, determined using a commercial Bicinchoninic acid assay (ThermoFisher Scientific) according to manufacturer’s instructions.

The expression of 111 proteins of interest was semi-quantitatively measured in pooled decidual lysates using the Mouse XL Cytokine Proteome Profiler Array (R&D Systems). Tissue lysates were thawed and clarified by centrifugation, and total protein concentration was measured using a commercial BCA assay as above. Samples were pooled by maternal diet and the sex of the conceptus from which the matched decidual tissue was isolated (CON = 4 – 6/sex, HFHS = 6/sex) and a total of 100 μg of protein from each pool (containing equal contributions by amount of protein) were incubated with each membrane, according to the manufacturer’s instructions. Membranes were developed and imaged simultaneously by chemiluminescence on a Bio-Rad Chemidoc set to high sensitivity. Densitometric signal intensity was measured in duplicate using the ‘Volume’ tool with global background subtraction in ImageLab software (Bio-Rad, https://www.bio-rad.com/en-ca/product/image-lab-software) Semi-quantitative differences in protein expression were measured by comparing the fold difference of HFHS samples to CON samples within each sex. A complete list of analytes, membrane coordinates, and their signal intensities for each membrane are available in Supplementary Table 5.

### Statistics

All statistical analyses in this study were performed in R (https://www.R-project.org). One litter or dam was considered a biological replicate, with one male and female embryo/implantation site from each litter used where data were stratified by sex. Pairwise comparisons were made using univariate linear models with maternal diet as the dependent variable, or Mann-Whitney U-test where the data were discrete values (e.g., litter size) or non-normally distributed (determined by D’Agostino-Pearson K2 test). For experiments with repeated measures, data were analyzed using mixed effects linear models using the lmerTest package (74) (version 3.1-3, SCR_015656) in R (v.4.2.3) with maternal diet and time (maternal weight gain and OGTT) or diet and sex (histological analyses) as fixed effects, and subject (dam) or sample (conceptus) as a random effect, respectively. The main effects of diet and time/embryonic sex were measured in these models with a two-way ANOVA using Type 3 sum of squares and Satterthwaite’s method for calculating denominator degrees of freedom. Where significant main effects of interactions were present, pairwise post hoc comparisons were made between the estimated marginal means using the emmeans package (75) (v.1.8.5, RRID SCR_018734). For all analyses, p < 0.05 was used as a nominal threshold for determining statistical significance.

### Study approval

All animal procedures for this study were approved by the McMaster University Animal Research Ethics Board (Animal Utilization Protocol 20-07-27), in accordance with the guidelines of the Canadian Council on Animal Care.

### Data availability

Data for this study will be made available upon reasonable request to the corresponding author.

## RESULTS

### Chronic high-fat, high-sucrose diet feeding induces maternal excess adiposity and dysglycemia

A timeline of preconception dietary intervention, metabolic testing, mating, and sample collection are outlined in Figure 1a. Preconception body weights were significantly higher in HFHS females compared to CON females from eight weeks of dietary intervention, and this difference was maintained into pregnancy (Figure 1b). Relative to their initial body weight at diet onset, HFHS females had gained twice as much weight as their CON counterparts by 12 weeks of dietary intervention and at the time they became pregnant (Figure 1c). HFHS dams did not exhibit major fertility impairments, with mating efficacy, litter sizes, and rates of embryonic resorption comparable to that of CON females (Table 1).

**Figure 1.**
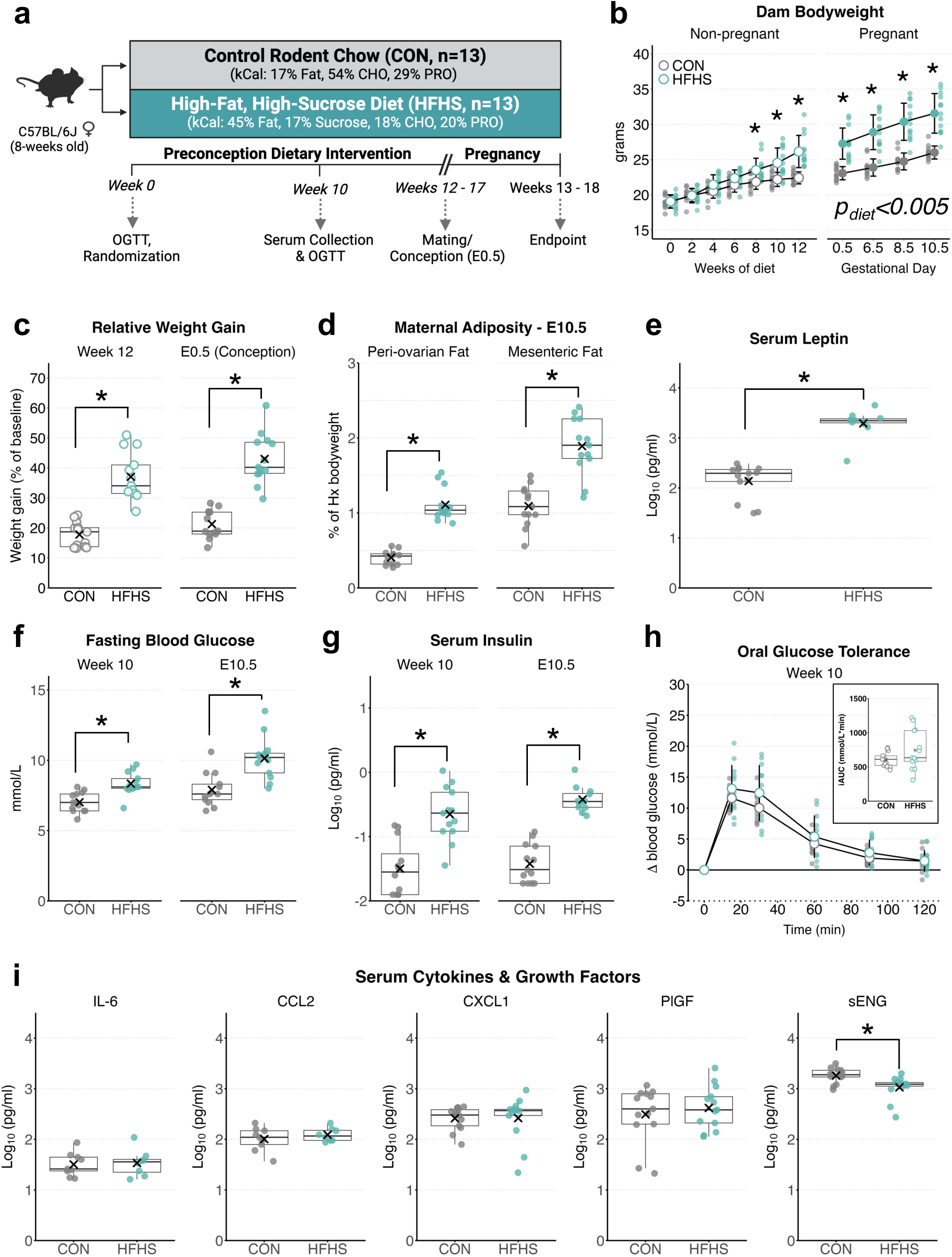
Murine model of diet-induced excess-adiposity. **(a)** Experimental timeline; pregnant dams were euthanized at embryonic day 10.5 (E10.5) following 12-17 weeks of preconception dietary intervention and tissues were collected for immunophenotypic, histological, and molecular analyses. **(b)** Preconception body weights during the first 12 weeks of dietary intervention. **(c)** Maternal weight gain relative to baseline body weight measured following 12 weeks of dietary intervention and at conception (E0.5). **(d)** Maternal gonadal and mesenteric adipose depot weights expressed relative to maternal weight following hysterectomy at E10.5. **(e)** Serum leptin levels at E10.5. **(f)** Six-hour fasting blood glucose and **(g)** serum insulin levels following 10 weeks of diet (preconception) and during pregnancy (E10.5). **(h)** Changes to blood glucose levels following oral glucose challenge (2g/kg body weight) at 10-weeks of dietary intervention; inset graph – area under the glucose curve. **(i)** Multiplex profiling of circulating inflammatory, and angiogenic mediators in maternal serum at E10.5 using multiplex magnetic bead immunoassay (from left – right, top row: Interleukin 6 (IL-6), chemokine C-C motif ligand 2 (CCL2), chemokine C-X-C-motif ligand 1 (CXCL1), Placental growth factor (PlGF), soluble Endoglin (sENG). Data for control (CON, n = 8 – 13) and high-fat, high-sucrose fed (HFHS n = 9 – 13) are presented as mean ± standard deviation (SD) underlaid with individual data points from females in each group in (b and h). Boxplots in (c – g, i) display IQR (box), and median (line), with whiskers indicating min/max values for each group within 1.5SD of the mean (denoted by ‘X). Hollow points indicate measures taken before conception, and solid points indicate measures taken during pregnancy. Pairwise comparisons were made using linear regression with maternal diet as the independent variable, and time-course data were analyzed using linear mixed-effects models with maternal diet and time as fixed effects, and subject (dam) as a random effect to nest repeated measures. Where the main effects of diet were statistically significant, post hoc pairwise post hoc comparisons of estimated marginal means were made between diet groups. * Indicates p<0.05 for pairwise comparisons.

**Table 1.** Measures of mating efficacy and reproductive outcomes at E10.5.

| Measure | CON (n=13) | HFHS (n=13) | *p*-value |
| --- | --- | --- | --- |
| Maternal age (weeks) | 23.2<br>(20.7 – 25.9) | 22.9<br>(20.9 – 24.7) | 0.53<sup>b</sup> |
| Pairings to 1<sup>st</sup> plug | 3.38<br>(1 – 8) | 3.62<br>(1 – 11) | 0.62<sup>a</sup> |
| Plugs to pregnancy | 1.31<br>(1 – 2) | 1.31<br>(1 – 2) | 1.00<sup>a</sup> |
| Total implantation sites | 8.77<br>(4 – 11) | 9.00<br>(4 – 10) | 0.55<sup>a</sup> |
| Resorption rate (%) | 7.37<br>(0 – 22.2) | 2.31<br>(0 – 20.0) | 0.16<sup>a</sup> |
| Viable conceptuses | 8.08<br>(4 – 11) | 8.77<br>(4 – 10) | 0.14<sup>a</sup> |
| Gravid uterine weight (g) | 1.17<br>(0.6 – 1.9) | 0.98<br>(0.3 – 1.6) | 0.18<sup>b</sup> |

Data are presented as mean with range (min – max). Data were analysed using <sup>a</sup>Mann-Whitney U-test or <sup>b</sup> univariate linear regression.

As expected, HFHS diet-induced weight gain was linked to increased adipose tissue accretion, including in visceral peri-ovarian and mesenteric depots (Figure 1d), and increased serum leptin levels at mid-gestation (E10.5; Figure 1e). Hyperglycemia and hyperinsulinemia present before conception were also pronounced in pregnant dams as well (Figure 1f and g). However, preconception dysglycemia was not associated with markedly impaired glucose disposal in HFHS females following an oral glucose challenge (Figure 1h), although they did remain hyperglycemic relative to CON females throughout the test (Supplementary Figure 2). Overall, the degree of excess adiposity and metabolic compromise we observed in HFHS-fed dams resembles the metabolic phenotype reported in pregnant women who are overweight or moderately obese (44, 45, 48, 76–81) and previous animal models of diet-induced obesity in pregnancy (82–85). In line with this, levels of stereotypical inflammatory mediators IL-6, CCL2, and CXCL1 were comparable between CON and HFHS dams, and TNF was undetectable in serum samples from both groups, consistent with previous reports in both humans and other animal models of diet-induced excess adiposity (48, 50, 60, 77, 83, 86) (Figure 1i). We also measured circulating levels of soluble Endoglin (sENG) and placenta growth factor (PlGF), which have been shown to increase and decrease in maternal serum in association with gestational hypertensive disorders, respectively (87–89). Interestingly, while PlGF levels were unaffected, sENG were considerably reduced in HFHS dams, indicating a lack of systemic anti-angiogenic shifts at mid-gestation (Figure 1i).

### Decidual innate lymphoid cell (dILC) and macrophage abundance are altered in HFHS pregnancies

To determine the impacts of this maternal obesogenic milieu on major decidual leukocyte populations during placental development, we performed tissue immunophenotyping of decidual ILC (dILC) and macrophage populations using multi-colour flow cytometry. (Figure 2a). Consistent with previous investigations, dILCs were predominantly comprised of pro-angiogenic CD49a^+^ EOMES^+^ tissue resident dNK cells (trNK) (∼75% of CD45^+^, Lin^-^, F4/80^-^, CD122^+^ cells), with smaller populations of CD49a^+^ EOMES^-^ type 1 ILCs (ILC1, ∼15%) and CD49a^-^ EOMES^int^ conventional-like NK cells (cNK, ∼10%) (53, 71, 90), the latter of which contribute to the initiation of spiral artery media destruction (25). Two decidual macrophage populations (CD45^+^, Lin^-^, CD122^low^, CD11b^hi^, Ly6C^low^, F4/80^hi^) were classified based on the presence (∼55%) or absence (∼45%) of MHC-II expression (67, 91) (Figure 2a).

**Figure 2.**
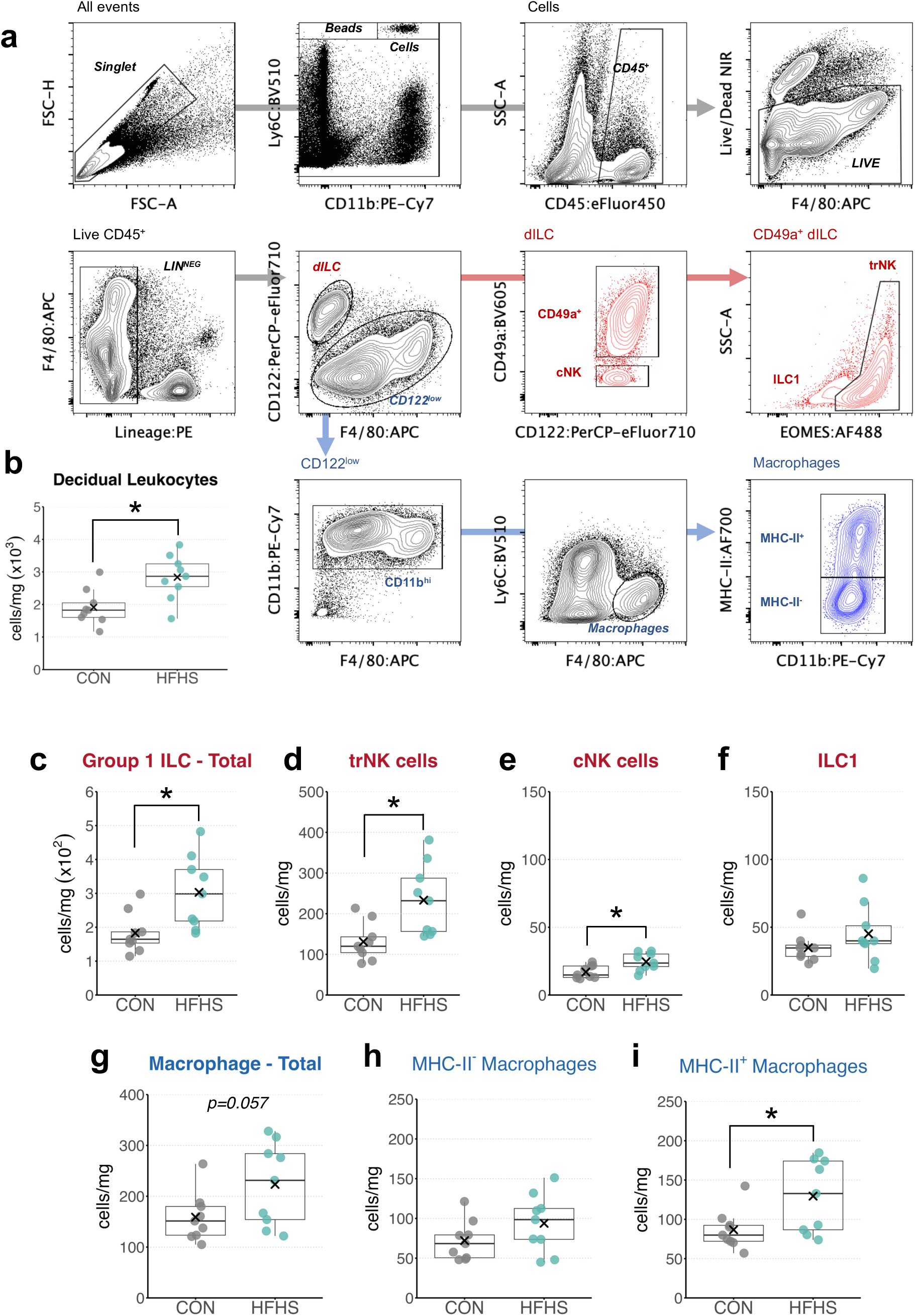
Maternal HFHS diet alters decidual immune cell abundance at mid-gestation. **(a)** Representative gating strategy to define innate lymphoid (dILC) and macrophage subsets in decidual tissue digests; arrows indicate gating hierarchy; lineage (LIN) markers – CD3e, CD19, Ly6G, and SiglecF. **(b)** Calibrated absolute counts of leukocytes (single, live, CD45^+^ events) per implantation site for CON (n=9) and HFHS (n=9) decidua. Each data point represents one measurement per pregnancy, boxplots display IQR (box) and median (center line) with whiskers depicting min and max values within 1.5 SD of the mean (denoted by X). Data were analysed by pairwise comparison using univariate linear models with maternal diet as the independent variable. * Indicates p<0.05. dILC = decidual innate lymphoid cell; cNK = conventional-like natural killer cell, trNK = tissue-resident (CD49a^+^) decidual natural killer cell; ILC1 = type 1 innate lymphoid cell.

Given the high vascular density of the decidua, circulating maternal cells contaminate the resident leukocyte pool in dissociated suspensions (6, 67). As macrophages and CD49a+ dILCs do not circulate at appreciable levels in blood (92, 93), quantities of these tissue-resident cells are more appropriately expressed according to their absolute abundance in decidual tissue, rather than a frequency of total leukocytes. Quantification of absolute cell counts revealed a near 50% increase in decidual leukocytes in samples from HFHS pregnancies (Figure 2b). Surprisingly, and in contrast to previous reports (53, 54), HFHS deciduae contained ∼65 % more dILCs per unit weight of decidua than CON, driven by increases in total numbers of both trNK (∼77% increase) and (to a lesser extent) cNK cells (∼45% increase) but not ILC1 (Figure 2c and d). Consequently, trNK made up a higher proportion of CD49a+ resident ILCs at the uteroplacental interface in HFHS pregnancies. This was accompanied by a marginal increase in total decidual macrophages in HFHS pregnancies (p=0.057, Figure 2f), attributable to a significant 50% increase in the number of MHC-II^+^ (p = 0.026) but not MHC-II^-^ cell counts (Figure 2g). The ratio of macrophage subsets was similar between diet groups (data not shown). These effects were similar regardless of whether cell counts were normalized to tissue weights or the number of implantation sites, except that the strength of the effect of diet was weaker for both cNK and MHC-II^+^ macrophages (each p=0.06) and the effect on total macrophage counts was abolished (p=0.15) when expressed in this manner (Supplementary Figure 3a and b). Strikingly, none of these differences in decidual leukocyte complement were captured when dNK and macrophage populations were expressed as a proportion of total leukocytes in decidual cell suspensions, as reported previously (53, 54). Instead, the sole difference between CON and HFHS pregnancies was a modest (∼17%) reduction in ILC1 cells. Overall, these findings demonstrate a previously unappreciated impact of diet-induced excess adiposity on the composition of decidual macrophages and ILCs at mid-gestation that is masked by concomitant increases in the abundance of multiple leukocyte populations when using frequency-based quantifications (Supplementary Figure 3c and d).

### Remodelling of decidual spiral arteries is not impacted in HFHS pregnancies

Given the changes we observed in innate immune cells and prior observations of reduced placental oxygenation and pathology suggestive of placental malperfusion in HFHS pregnancies (38, 64, 84, 86), we performed a histological examination of implantation sites to assess their overall structure and the progression of modifications to maternal spiral arteries at E10.5, a period of active immune-mediated remodelling (94). As data have suggested that susceptibility to pathologies associated with insufficient spiral artery remodelling may be sex-biased (95–97), we stratified our analyses based on the sex of each conceptus occupying each implantation site. A gross comparison of mid-sagittal sections from CON and HFHS pregnancies did not reveal any consistent abnormalities in tissue ultrastructure between groups (Figure 3a-b, and Supplementary Figure 4). Moreover, and in contrast to previous reports (51, 53), we did not find strong evidence of reduced arterial lumen size or vessel cross-sectional area between diet groups. There was a tendency for an increased wall-to-lumen area ratio in spiral arteries supplying HFHS female placentas specifically, but this did not reach statistical significance (main effect of diet, p=0.110; effect of interaction, p=0.106, Figure 3d). Still, we did observe that decidua of HFHS implantation sites tended to exhibit greater cellularity and appeared more edematous compared to those from CON pregnancies (Figure 3a and b), consistent with our finding of an overall increased leukocyte abundance in HFHS decidual cell suspensions, and potentially indicative of heightened local inflammation.

**Figure 3.**
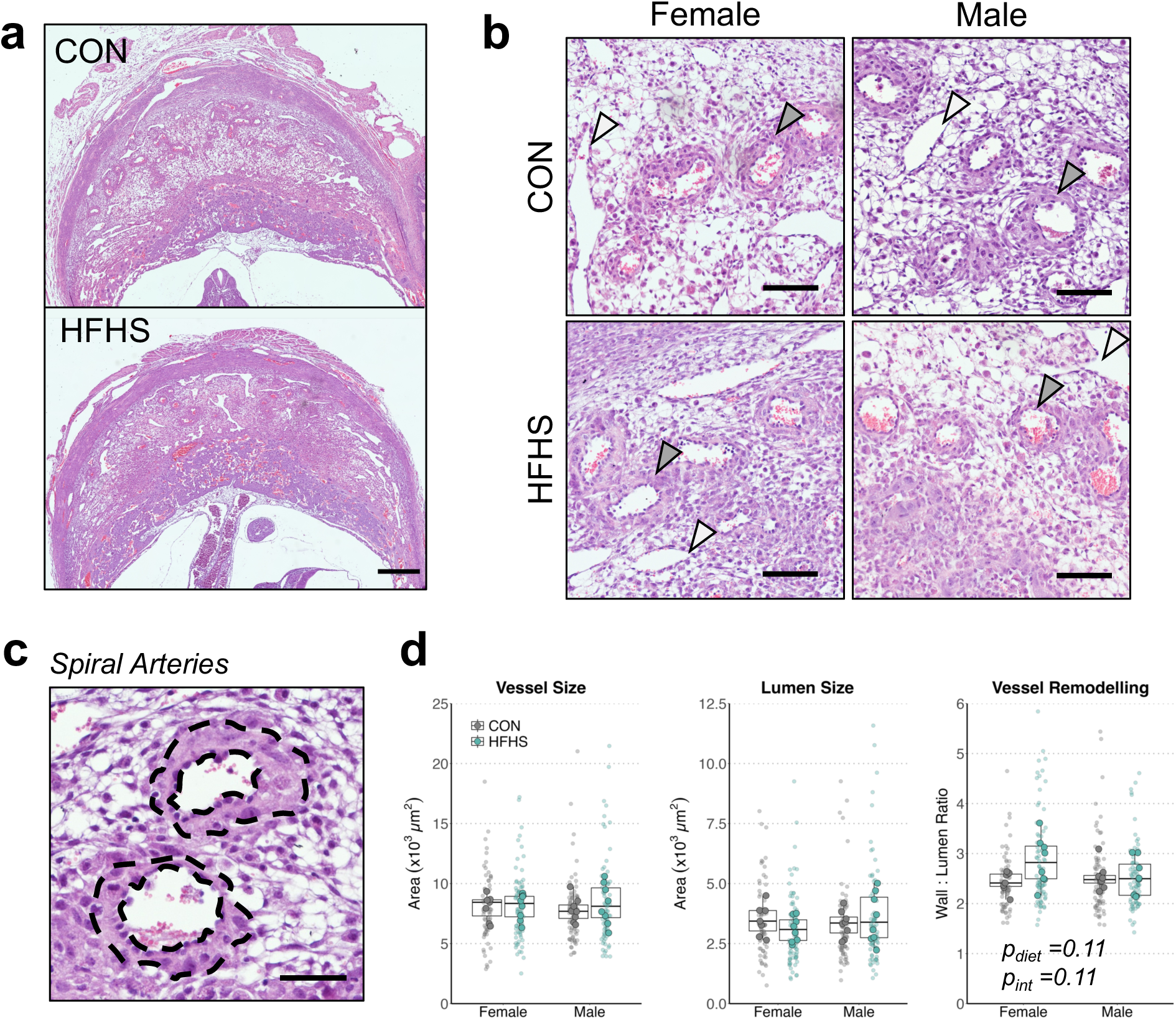
Maternal HFHS diet does not alter gross spiral artery morphology at mid-gestation. **(a)** Representative composite images of mid-sagittal E10.5 implantation site from CON and HFHS pregnancies; scale bar = 500 μm. **(b)** Representative fields of H&E-stained decidua from CON and HFHS pregnancies associated with female or male conceptuses. White arrows indicate decidual veins, identified by thin-walled endothelium, which were excluded from remodelling analyses. Grey arrows indicate examples of spiral artery cross sections that were included in analyses; tissues were imaged at 20× magnification; scale = 100 μm. **(c)** Representative zoning of spiral artery wall (outer outline) and lumen (inner outline) from H&E stained decidua, field taken from decidual tissue imaged at 20× magnification; scale = 50μm. **(d)** Spiral artery wall (left) and lumen (middle) area measurements, and their ratio (right) taken from CON (n = 7 – 8/sex) and HFHS (n = 8/sex). Data are presented with boxplots displaying IQR (box) and median (center line), with min and max values with 1.5 IQR (whiskers) calculated according to the mean values per sample. Small data points on each plot represent individual arterial measurements from all samples belonging to each group; larger data points represent the mean value of all observations per implantation site for each measure presented. All data were analysed using two-factor ANOVA of mixed-effects linear models, with diet and sex as fixed effects and sample ID as a random effect.

To more definitively resolve the impacts of the obesogenic environment on metrics of spiral artery remodelling, immunofluorescent staining of implantation sites for alpha-smooth muscle actin (α-SMA), Vimentin, and Ki-67 was performed to compare the degree of residual contouring of spiral arteries by vascular smooth muscle and angiogenesis within the vessel intima (Figure 4a-b). Consistent with the morphometric analyses, the perimeter of the endothelial layer of spiral arteries was similar between diet groups, as was the extent of vascular smooth muscle removal from the arterial media (Figure 4c). Ki-67 positivity among endothelial cells of spiral arteries was also comparable between CON and HFHS implantation sites (Figure 4c). The number of arterial lumens per decidual cross-section was also not significantly different between groups (Figure 4c). Overall, these findings demonstrated that the progression of vascular media destruction and angiogenesis in the decidual segments of the spiral arteries was not remarkably impacted by maternal diet, despite pronounced differences in the abundance of NK cells and macrophages implicated in their remodelling.

**Figure 4.**
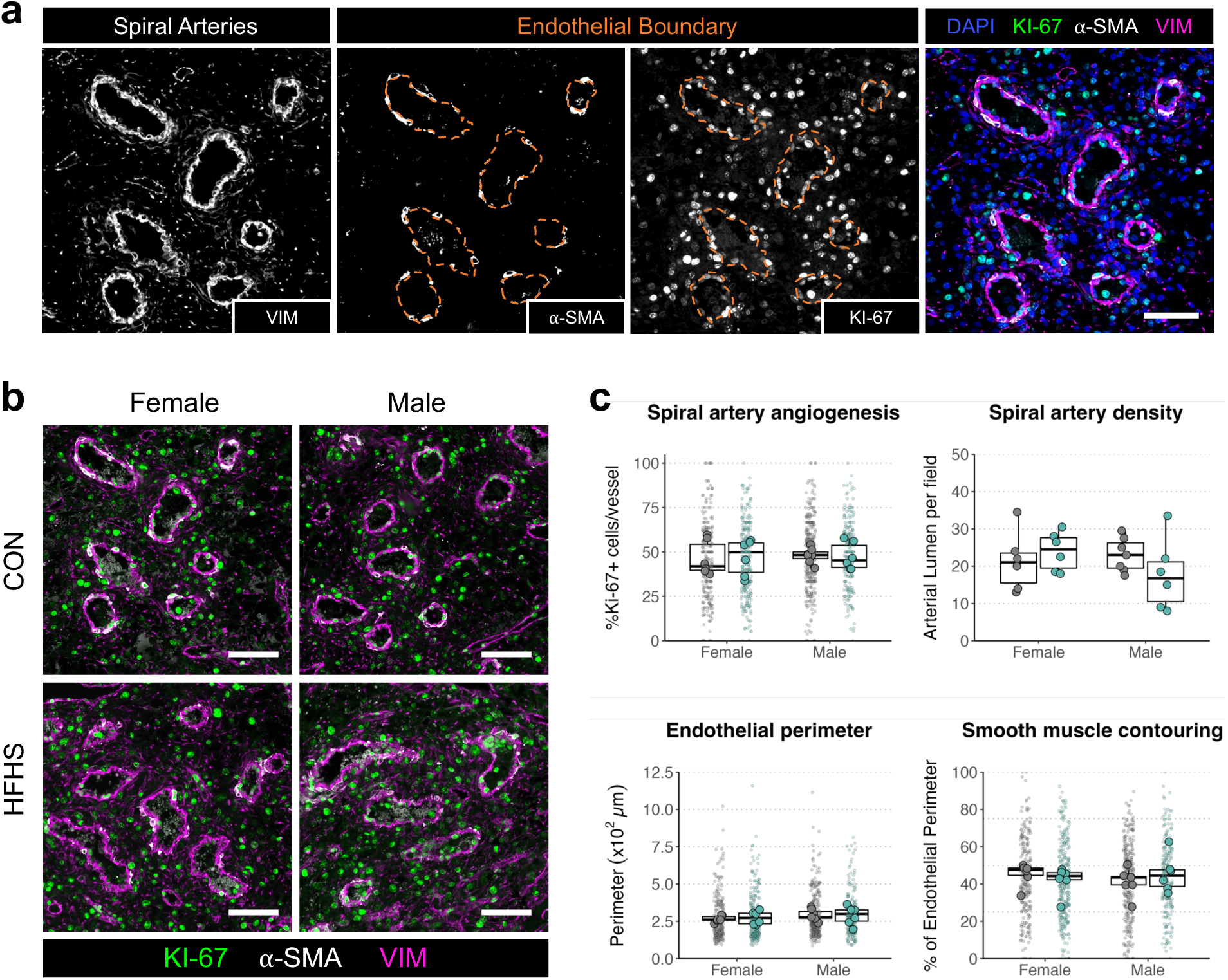
Maternal HFHS diet does not alter decidual spiral artery angiogenesis or smooth muscle destruction at mid-gestation. **(a)** Representative immunofluorescence staining of E10.5 decidua for Vimentin (VIM, left), smooth muscle alpha-actin (α-SMA, middle-left), KI-67 (middle-right), and their co-localization (right); hashed outlines = boundary of the endothelial layer of each spiral artery; fields taken from whole tissue scans imaged at 20× magnification; scale = 100μm. **(f)** Representative spiral arteries immunostained for VIM (magenta), α-SMA (white), and Ki-67 (green) from CON and HFHS decidua associated with female or male conceptuses; Fields images at 20× magnification; scale = 100μm. **(g)** Indices of spiral artery remodelling quantified in decidua from CON and HFHS dams associated with male or female conceptuses: endothelial proliferation was measured as the percent of total DAPI-stained endothelial cell nuclei (luminal VIM^+^ α-SMA^-^) per spiral artery (top-left); the total number of arterial lumen per mid-sagittal section of each embryonic implantation site was measured and averaged across each implantation site (top right); the perimeter of each vessel intima (endothelial layer) was measured across duplicate mid-sagittal sections of embryonic implantation sites from CON and HFHS decidua (bottom left); intactness of the vascular smooth muscle layer was measured by calculating the fraction of each endothelial perimeter contoured by intact α-SMA staining (bottom right). Data are presented with boxplots displaying IQR (box) and median (center line), with min and max values with 1.5 IQR (whiskers) calculated according to the mean values per sample. Small data points on each plot represent individual arterial measurements from all samples belonging to each group; larger data points represent the mean value of all observations per implantation site for each measure presented. All data were analysed using two-factor ANOVA of mixed-effects linear models, with diet and sex as fixed effects and sample ID as a random effect, except for counts of arteries/field in (g), which were analysed by regular two-way ANOVA.

### Expression of inflammatory and angiogenic mediators is enhanced in HFHS decidual tissue

To explore whether physiological shifts imparted by HFHS diet-induced excess adiposity impacted facets of the decidual environment outside of spiral artery remodelling, we compared protein expression levels of signalling factors with established roles in inflammation, angiogenesis, and vascular function between CON and HFHS decidual tissue using both targeted and exploratory approaches. We first profiled a panel of 12 targets consisting of cytokines, chemokines, and growth factors with known roles in both decidual angiogenesis, inflammation, and placental development, many of which are expressed by dNK and macrophage populations. This revealed profound increases in protein expression of both vascular endothelial growth factor isoform A (VEGF-A) and placenta growth factor (PlGF) with six- and nine-fold increases in their mean expression in HFHS decidual tissues, respectively (Figure 5a). HFHS decidua also contained significantly elevated levels of leptin and proinflammatory tumor necrosis factor (TNF). CXC-motif ligand 1 (CXCL1), a chemokine with known roles in neutrophil recruitment and endothelial migration (98, 99) and known to be induced by TNF (100), also showed a tendency for increased expression (p = 0.072). Interestingly, levels of hepatocyte growth factor (HGF), were reduced in HFHS decidua. HGF is expressed by decidual stromal cells (101, 102) and trNK cells (103, 104) in humans and mice, with putative roles in mediating trophoblast proliferation (101), alternative macrophage polarization (105), and angiogenesis (106).

**Figure 5.**
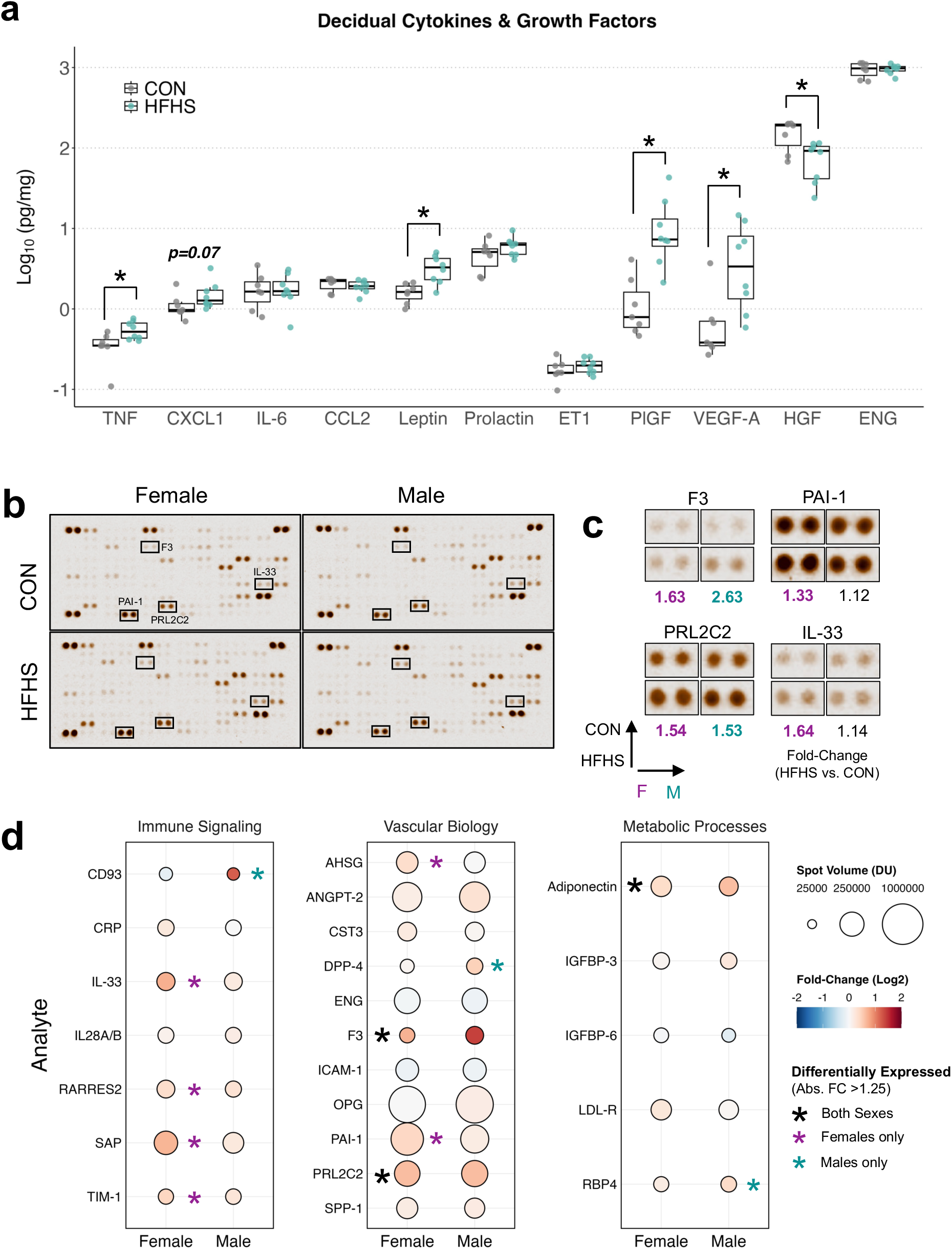
HFHS diet alters decidual protein expression of inflammatory and angiogenic mediators at E10.5. **(a)** Relative protein levels (pg of analyte relative to mg of total protein in sample) of cytokines, chemokines, growth factors, and mediators of vascular function in decidual tissue lysates from CON (n=7) and HFHS (n=8) pregnancies at E10.5. **(b)** Representative image of protein array membranes of pooled decidual lysates from CON (n=4-6/sex) and HFHS (n=6/sex) decidua. **(c)** Insets identified in (b) depicting representative paired spots for single analytes with fold-change (FC) ≥1.25 (bolded text) for at least one sex (females (F) = magenta, males (M) = teal) between CON and HFHS decidua. **(d)** Dot plot of all 23 reliably detected target analytes in the protein array. Target analytes were grouped according to known functions and gene ontology (GO) annotations into three categories – immune signalling, vascular biology, and metabolic processes. Dot colour corresponds to the Log_2_(Fold-Change) (HFHS versus CON) for comparisons within each sex. Dot size corresponds to the absolute spot volume (protein expression) at a given exposure in densitometric units (DU). Asterisks indicate target analytes considered to be differentially expressed in both sexes in the same (black) or opposite (grey) direction, or exclusively in females (magenta) or males (teal). Data in (a) are presented with box plots depicting the IQR (box), median (center line), and the min/max values within 1.5 units of the IQR (whiskers). * Indicates p<0.05 for pairwise comparisons. Data in (a) were analyzed by pairwise comparison with Welch’s t-test or Mann-Whitney U-test. A list of all analytes and corresponding coordinates in (b) are listed in Supplementary Table S4.4. F3 = Factor 3 (Tissue Factor), PAI-1 = Plasminogen activator inhibitor 1, PRL2C2 = Prolactin family 2, subfamily c, member 2 (Proliferin), IL-33 = Interleukin 33.

Several of the factors elevated within the decidua of HFHS dams have roles in both the normal progression of spiral artery remodelling (3, 14, 94, 107–109), as well as endothelial activation/ inflammation (61, 110–112), and are regulated tissue oxygenation (113, 114) To help contextualize the pattern of changes observed in decidual immune cell abundance and cytokine levels, we next profiled a broader set of signalling factors altered in HFHS tissues. Decidual lysates were pooled by diet and sex and profiled using a membrane protein array. Out of 111 analytes included in the array (Supplementary Table 5), 23 were detected at a level that allowed for reliable comparison of expression differences (Figure 5b). Comparing expression between CON and HFHS tissues within sex, a nominal cut-off of ±1.25-fold change in HFHS relative to control was used to identify differentially expressed targets considered to be biologically meaningful (53). Based on these criteria, 12 proteins showed differential expression in HFHS decidua, all of which were increased relative to control. Adiponectin, Tissue Factor (F3), and Proliferin (PRL2C2) were increased in both HFHS decidua of males and females; Interleukin-33 (IL-33), Retinoic acid receptor responder protein 2 (RARRES2/Chemerin), T-cell immunoglobulin and mucin domain 1 (TIM-1), Alpha-2-HS-glycoprotein (AHSG/Fetuin A), Plasminogen activator inhibitor-1 (PAI-1), and Serum amyloid P component (SAP) showed more than a 1.25-fold increase but in HFHS females only; and CD93, Dipeptidyl peptidase-4 (DPP4), and Retinol binding protein 4 (RBP4) showed ≥1.25-fold increase in HFHS males only (Figure 5c). Among the analytes identified, F3 showed the greatest increase, with 2.6-fold and 1.6-fold increases in HFHS decidua associated with male and female conceptuses, respectively. Several of the factors identified to be differentially expressed have known roles in initiating thrombosis (F3) (115) or inhibiting fibrinolysis (PAI-1) (116), are induced in response to tissue injury (SAP, IL-33) (117, 118), or promote endothelial cell migration and neovascularization (PRL2C2) (119). Collectively, these data suggest that maternal HFHS diet-induced excess adiposity perturbs decidual vascular and immune cell homeostasis during the active phase of uterine remodelling and placental development. These data reinforce that changes in pathways involved in vascular inflammation, coagulation, and/or upstream changes to uteroplacental perfusion, rather than pronounced impairments to initiation of immune-mediated spiral artery remodelling, likely contribute to placental inflammation and compromised maternal perfusion characteristic of pregnancies impacted by maternal disorders.

## DISCUSSION

Our previous work and that of others have indicated signs of placental malperfusion, hypoxia, and inflammation in pregnancies complicated by diet-induced excess adiposity (55, 83, 84, 120–123). Data from human pregnancies has suggested that maternal obesity may impede the early remodelling of spiral arteries by way of reduced dNK cell abundance, potentiating these adverse outcomes (51, 124). Here, we explored whether a similar mechanism of arterial remodelling impairments associated with shifts in the decidual immune environment at mid-gestation preceded signs of placental dysfunction observed in HFHS diet-fed dams later in pregnancy (125, 126).

Chronic preconception HFHS diet consumption induced excess maternal adiposity, hyperglycemia, hyperinsulinemia, and hyperleptinemia – all features observed in human pregnancies complicated by obesity (51, 78, 127, 128). Contrary to our hypothesis, we did not observe any penetrant changes to common metrics of spiral artery remodelling in HFHS pregnancies at mid-gestation, including vascular smooth muscle loss, endothelial proliferation, and vessel size. Moreover, populations of dNK cells were not decreased, but instead increased at mid-gestation, alongside increases in MHC-II^+^ macrophage numbers. Decidual tissue of HFHS dams also had higher protein expression of several pleiotropic growth factors and cytokines many of which have overlapping roles in both the normal progression of placentation, as well as pathologic inflammation including pronounced increases in leptin (129, 130), VEGF-A (131, 132), PlGF (111, 133), and F3 (115, 134), as well as TNF(3, 94), IL-33 (118, 135), PAI-1 (136, 137), and CXCL1 (100, 138) to a smaller extent. While the exact implications of these changes remain to be determined, the considerable shifts observed in decidual immune cell composition, inflammatory mediators and growth factors suggest that maternal HFHS intake disturbs normal decidual immune and vascular homeostasis in early pregnancy, which likely affects the course of placental development and function later in gestation.

The dNK subsets we find to be increased in HFHS decidua serve distinct roles in early placentation in mice. Principally, cNK cells produce high levels of IFN-γ, a cytokine essential for initiating spiral artery transformation as well as the maturation of trNK (103, 139, 140), which promote decidual angiogenesis and fetoplacental development through the regulation of trophoblast invasion (3, 23), production of growth-promoting factors (12), and pro-angiogenic stimuli (103, 107, 141). This includes the production of VEGF-A and PlGF, both of which we observed to be markedly higher in HFHS decidual tissues. The increases in the expression of VEGF-A and PlGF in these tissues may reflect the increased abundance of trNK cells which express them (107). However, the accumulation of cNK and trNK cells as well as differences in magnitude compared to increases in VEGF-A and PlGF levels (1.5-fold vs. 6- and 9-fold), suggest that normal dNK cell function may be altered.

In line with this, a recent study in mice similar to ours has demonstrated that bulk dNK cells from HFHS pregnancies have reduced *Vegfa* expression at E10.5 (53). Similarly, maternal obesity has been associated with reduced in vitro VEGF-A secretion in first-trimester decidual leukocytes (52) which predominantly consist of dNK cells, pointing to a defect in the pro-angiogenic functions of these cells (52). The angiogenic effector functions of dNK cells are linked to a process of phenotypic maturation (11, 107, 142, 143) and may depend in part on IFN-γ signalling between cNK and trNK subsets (103, 139). In alymphoid mice engrafted with IFN-γRa^-/-^ bone marrow, spiral artery remodelling is comparable to decidua of wildtype mice as IFN-γ signalling to other decidual cells remain intact, but dNK cells fail to mature and accumulate excessively (139). This scenario resembles the patterns we observe here, suggesting that the ability of these cells to properly mature could be impaired or delayed and that perhaps compensatory increases are needed to affect the same level of vascular remodelling in HFHS pregnancies. Production of IFN-γ by peripheral blood NK (pbNK) cells is reduced in obese individuals, which may be mediated by chronic exposure to saturated fatty acids (SFA) (144), (which are elevated in our HFHS diet) and leptin (145), (which we show to be elevated in HFHS decidual lysates). SFA and leptin may have similar effects on decidual cNK populations and their ability to induce trNK maturation, but this remains to be determined. Furthermore, decidual levels of HGF, a cytokine produced by trNK in humans and mice (103, 104), is reduced in our HFHS decidua. HGF produced by dNK cells in humans has been suggested to promote uterine angiogenesis and trophoblast invasion (101, 106) and is reduced in dNK cell culture supernatants from pregnancies with high uterine artery resistance indices – a proxy measure of poor placentation (104). This evidence further supports our hypothesis that dNK cell function may be adversely impacted by maternal HFHS intake in our model.

In addition to their role in the remodelling of spiral arteries, dNK cells also facilitate the expansion and pruning of microvascular networks supplying the decidua itself (11). Experimental depletion of dNK cells in rodent models leads to blunted decidual angiogenesis and hypoxia (21, 146), activating conserved adaptive responses that promote compensatory trophoblast invasion to achieve adequate placental perfusion (21, 146–148), Although we have yet to perform detailed histological examinations of CON and HFHS placentas at this stage, we found that HFHS decidual samples had higher expression of Proliferin (PRL2C2), a factor specifically secreted by invasive spiral artery-associated trophoblast giant cells (TGCs) in mice (119, 149). Previous work has also demonstrated precocious interstitial and endovascular trophoblast invasion in the decidua of chronic HFHS diet-fed rats, coincident with uteroplacental hypoxia (55, 64). Factors that facilitate this compensatory remodelling response include VEGF-A, IL-33, and MMP-9 (150), which we and others (55) have now shown to be increased in HFHS decidual tissue. Thus, the changes we observe may indicate that adaptive responses to intrauterine hypoxia arising from dNK cell-related angiogenic deficits are occurring and affect downstream changes in trophoblast lineage differentiation and placental development.

Consistent with this hypothesis, Ncr1^-/-^ mice, which exhibit delays in dNK cell maturation and decidual vascularization, also show elevated VEGF-A expression in the decidual stroma at mid-gestation. This may reflect yet another compensatory angiogenic response to hypoxia (151). Similar decidual overexpression of VEGF-A has been demonstrated in decidua of preeclamptic (PE) pregnancies (152). Additionally, decidual hypoxia in PE cases has been directly linked to increases in leptin expression (113, 153), consistent with our data. Both leptin (154–157) and VEGF-A (158–162) have been shown to facilitate multiple aspects of trophoblast survival and invasion, and recent work has shown that transgenic overexpression of VEGF-A by decidual cells increases the numbers of TGCs in mice and also increases the size of maternal blood sinusoids in the placental labyrinth (162). Expression of the leptin receptor has also been recently demonstrated on sinusoidal TGCs which line these spaces, arising around E10.5, suggesting a role in their morphogenesis (129). We cannot differentiate between increases in decidual leptin production and elevated maternal circulating leptin levels in HFHS dams, thus further studies localizing the expression of these cytokines within decidual tissue are needed. While the cellular and molecular changes we demonstrate are suggestive of adaptations to dNK cell-related functional deficits, a more comprehensive examination of decidual vascular network structure, and dNK cell maturation throughout early placental development are needed to substantiate this.

Decidual hypoxia can also result from inflammation (163). The accumulation of dNK cells we observe may reflect a part of an inflammatory response. While dNK cells are usually poorly cytotoxic in normal pregnancies, it has been demonstrated that inflammatory challenges, including LPS, can lead to dNK cell expansion and activation, promoting their degranulation and release of cytolytic effectors and proinflammatory cytokines. This then leads to placental injury and potentially pregnancy loss (16, 164). Previous work in humans has shown increased basal and stimulated dNK cell degranulation in isolated first-trimester dNK cells from obese pregnancies, along with moderate increases in dNK cell expression of TNF, and pronounced increases in their secretion of PlGF (52), similar to the cytokine profile we show in HFHS decidual tissue. This may suggest greater dNK cell activation in these pregnancies, potentially in response to inflammatory cascades activated by bacterial endotoxin, which is increased in circulation (84) and at the interface (126) in high-fat diet murine models.

In keeping with the notion of a pro-inflammatory environment, we found that total decidual leukocyte numbers were increased in HFHS decidua, including increased MHC-II^+^ macrophages, elevated TNF levels, increased tissue factor (F3) expression, and upregulation of mediators involved in acute phase responses and coagulation (PAI-1, SAP). Tissue factor (F3) expression is potently induced by LPS stimulation (165, 166) and promotes activation of coagulation cascades (115). PAI-1 is an inhibitor of fibrinolysis and is induced by TNF (167). SAP is an acute-phase protein that binds pathogen-associated moieties like LPS (168) and promotes opsonization, complement activation, and phagocytic clearance (169). In addition to its synergistic role in angiogenesis, PlGF can also contribute to the induction of T_h_17 responses against extracellular pathogens (170), involving the production of TNF and CXCL1(171–174), which tended to be elevated in HFHS decidual tissues. Pathological T_h_17 skewing has been implicated in pregnancy disorders including spontaneous pregnancy loss and PE (18, 175). Together these findings suggest that a maternal HFHS diet disturbs the inflammatory balance at the maternal-fetal interface and leads to apparent procoagulant shifts during early placentation.

Despite these changes in immune homeostasis, we did not observe overt signs of placental or decidual developmental defects, nor did we see increased pregnancy loss at mid-gestation or later gestational time points (Chapter 3). This may suggest that these inflammatory challenges are effectively buffered. While increased MHC-II expression is typically associated with macrophage activation, inflammation, and promotion of effector T cell responses (176–178), decidual macrophages also bear several immune checkpoint ligands that inhibit effector T cell functions (4, 6, 14). Increases in human-term decidual HLA-DR^+^ macrophage frequency with obesity have been associated with reduced numbers of effector CD4^+^ and CD8^+^ T cells and increased frequencies of T_reg_ cells (17). Consistent with this, increased frequencies of T_reg_ cells in uterine draining lymph nodes during pregnancy have been shown in a mouse model of chronic preconception high-fat diet feeding similar to our own (179). We also observed that HFHS decidual tissue had higher expression of IL-33, a tolerogenic cytokine produced by type 2 uterine ILCs which buffers against endotoxin-induced pregnancy loss through inhibition of NK cytotoxicity and reprogramming of anti-inflammatory macrophages (180–182). Furthermore, we show increased expression of decidual adiponectin in HFHS tissues; adiponectin has known anti-coagulant, anti-inflammatory, and vasoprotective impacts (183–185). Thus, while the observed immune cell abundance might seem pathological, it could equally reflect an anti-inflammatory, pregnancy-sustaining response. In line with this, systemic vascular inflammation induced by intraperitoneal LPS administration has been shown to activate decidual IL-10 production to prevent preterm birth in mice (186).

It is worth noting that some of these molecular changes appear more frequently associated with female embryos, suggesting that conceptus sex could modify immune responses and pregnancy outcomes in metabolically compromised pregnancies, as previously described (64, 95, 187). Indeed, there was a notable tendency for altered arterial morphology specifically in HFHS females as well. Collectively these data point towards a complex inflammatory challenge in HFHS pregnancies that is modified in part by conceptus sex. While these inflammatory challenges may be effectively buffered, they may still have undesirable effects on downstream uteroplacental perfusion, placental development, and developmental programming in offspring.

Our finding that dNK cell numbers are increased in HFHS pregnancies lies in contrast to previous studies showing reduced dNK cell frequencies in first-trimester decidua of obese individuals (51), and unchanged frequencies in mice with diet-induced excess adiposity (53, 54). A few differences may account for these discrepancies. First, we have demonstrated that compositional differences in decidual macrophage and ILC populations are not apparent when expressed as a relative frequency, as has been exclusively reported in other human and rodent obesity studies (51, 53, 54). Likely, concomitant increases in several leukocyte populations under inflammatory conditions (as we see here) may obscure differences in absolute abundance. We believe our quantification of absolute numbers of cells more accurately reflects changes in decidual immune cell composition. Second, differences may be reflective of the gestational stage. Perdu et al. identified deficiencies in NK cell abundance in the early first trimester (51) (∼6 – 10 weeks of gestation), before the onset of placental perfusion, which would be akin to the E6.5 – E9.5 window in mice (4, 42, 188). How phenotypic and compositional changes in decidual leukocyte populations driven by metabolic dysfunction vary at different stages of pregnancy remains unaddressed in both humans and animal models, and should be investigated further. Finally, it is notable that the HFHS dams in our study do not exhibit signs of elevated soluble inflammatory markers, in contrast to what was observed in previous decidual immunophenotyping studies in humans and mice (51, 53). No one model can capture the entire spectrum of metabolic dysfunction and inflammatory sequelae in pregnancies complicated by excess adiposity, and it is possible that more severe metabolic dysfunction and associated inflammation manifest different consequences for dNK cell abundance and adverse pregnancy outcomes, including increased rates of embryonic demise seen in some obesogenic diet manipulations (53, 64, 189), but not in others (83, 84, 179, 187, 190).

We recognize that our study is not without limitations. Our evidence of increased immune cell abundance in HFHS decidua is limited to quantification using flow cytometry. While substantial upregulation of transcripts encoding leukocyte recruiting chemokines in the peri-implantation window has been observed in diet-induced obese rodents (50), enzymatic dissociation to isolate tissue-resident immune cells has known biases in dampening or enhancing differences in immune cell subsets (5). Paired immunohistochemical analyses are needed to confirm our findings, alongside detailed histopathologic comparisons between CON and HFHS uteroplacental tissues. Our immunophenotyping analyses are also limited in their ability to infer the functional capacity of these cells, and how the full complement of other resident decidual immune cell populations is affected, including T cells, dendritic cells, and granulocytes, remains to be examined. While our bulk decidual protein data support a model of altered immune cell abundance and function, future work employing omics-based methods at the single cell level would help to resolve how immune cell function and crosstalk with other cells, including trophoblast, endothelial cells, and stromal cells are impacted in metabolically complicated pregnancies.

In conclusion, we offer convincing evidence of disturbed decidual immune cells and vascular homeostasis early in placental development in pregnancies complicated by maternal HFHS diet exposure. Our results lend support to emerging notions from both human and rodent studies that impairments in spiral artery transformation are not ubiquitous among pregnancies impacted by excess adiposity, but are likely restricted to those pregnancies in which severe obstetrical complications arise (i.e., pregnancy loss, early-onset preeclampsia, or stillbirth) (124, 191). Still, local decidual vasculopathy and inflammation may contribute to the impedance of normal placental perfusion and tissue injury (38, 39, 121), which may precipitate adverse outcomes proximal to term gestation (i.e., late-onset preeclampsia) and/or participate in priming of chronic conditions in offspring postnatally (22, 192–195). We propose that enhancement of canonical inflammatory and angiogenic mediators may still be conducive to normal spiral artery remodelling while disturbing local coagulant balance, contributing to malperfusion, hypoxia, and adaptive changes in tissue-resident immune populations. Differing magnitudes in these disturbances may dictate the severity of later outcomes for fetal and placental development. Our findings demonstrate that processes involved in placentation are disturbed from early stages of pregnancy and highlight the need for a better understanding of their contributions to intrauterine adversity in pregnancies impacted by overweight and obesity.

## AUTHOR CONTRIBUTIONS

CJB, DMEB, and DMS conceived the study. DMS and AGS acquired funding. CJB, EY, TAR, and PAJ collected data. EY, TAR, PAJ, CE, JB, and AAA provided technical assistance. CJB performed all formal analyses and data visualization. CJB, AAA, AGS, DMEB, and DMS contributed to data interpretation. CJB drafted the manuscript. CJB and DMS edited the manuscript. DMS provided supervision. DMEB and DMS provided resources for this study. All authors have read and approved the manuscript in its final form.

## ACKNOWLEDGEMENTS

The authors thank Dr. Allison Kennedy, Dr. Jessica Breznik, and Erica DeJong for their assistance with establishing protocols for decidual immunophenotyping. AAA, DMEB, and DMS are supported by the Canada Research Chairs Program. CJB was supported by a Fredrick Banting and Charles Best Doctoral Award from the Canadian Institutes of Health Research. This work was supported by a Project Grant awarded to DMS and AGS by the Canadian Institutes of Health Research.

## SUPPLEMENTARY MATERIALS

**Supplementary Table 1.**
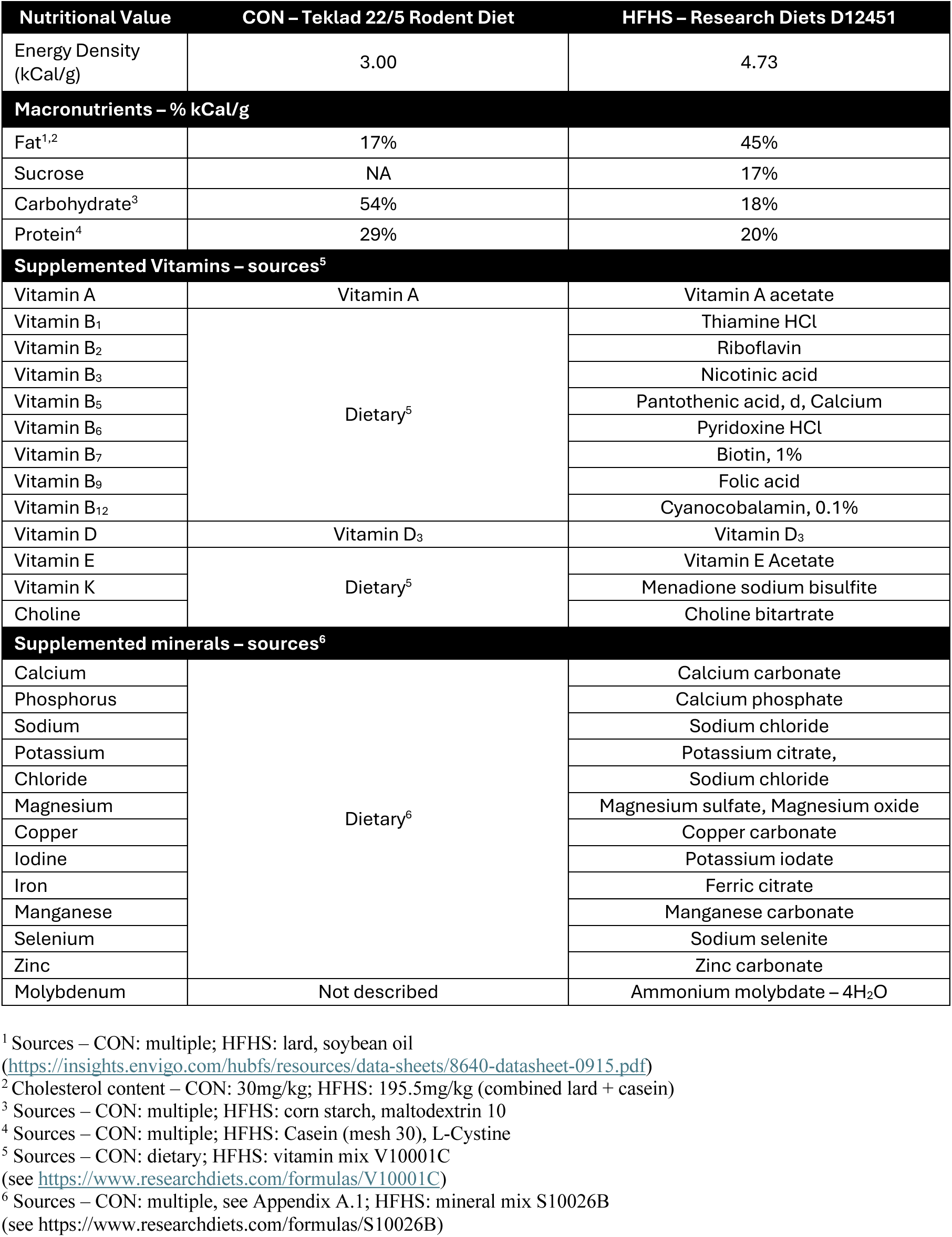
Nutrient composition comparison of experimental diets.

**Supplementary Table 2.** Fluorophore-conjugated antibodies used for decidual immunophenotyping.

| Marker | Fluorophore | Clone | Vendor – cat. No | Working dilution |
| --- | --- | --- | --- | --- |
| **Surface stain** |  |  |  |  |
| CD45 | eFluor450 | 30-F11 | TFS - 48-0451-82 | 1:200 |
| CD3e | PE | 145-2C11 | TFS - 12-0031-82 | 1:200 |
| CD19 | PE | 1D3 | TFS - 12-0193-82 | 1:200 |
| Ly6G | PE | 1A8-Ly6g | TFS - 12-9668-82 | 1:200 |
| SiglecF | PE | E50-2440 | BD - 552126 | 1:50 |
| CD11b | PE-Cy7 | M1/70 | TFS - 25-0112-82 | 1:100 |
| CD122 | PerCP-eFluor710 | TM-β1 | TFS - 46-1222-82 | 1:100 |
| Ly6C | BV510 | HK1.4 | BL - 128033 | 1:100 |
| F4/80 | APC | BM8 | TFS - 17-4801-82 | 1:100 |
| MHC-II (IA/IE) | AF700 | M5/114.15.2 | TFS - 56-5321-82 | 1:200 |
| CD49a | BV605 | Ha31/8 | BD - 740375 | 1:100 |
| Fixable Live/Dead-NIR | (APC-Cy7) | NA | TFS - L10119 | 1:250 |
| **Intracellular stain** |  |  |  |  |
| EOMES | AF488 | Dan11mag | TFS - 53-4875-82 | 1:100 |
| **Isotype control antibodies** |  |  |  |  |
| Rat IgG2b | PerCPeFluor710 | eB145/10H5 | TFS - 46403180 | 1:100 |
| Rat IgG2b | PE-Cy7 | RTK4530 | BL – 400617 | 1:100 |
| Rat IgG2a | APC | RTK2578 | BL - 400511 | 1:100 |
| Rat IgG2a | AF488 | eBR2a | TFS - 53-4321-80 | 1:100 |
| Rat IgG2b | AF700 | eB149/10H5 | TFS - 56-4031-80 | 1:200 |

TFS = ThermoFisher Scientific BD = BD Biosciences BL = Biolegend BV = Brilliant Violet AF. = AlexaFluor () = equivalent emission spectra

**Supplementary Table 3.**
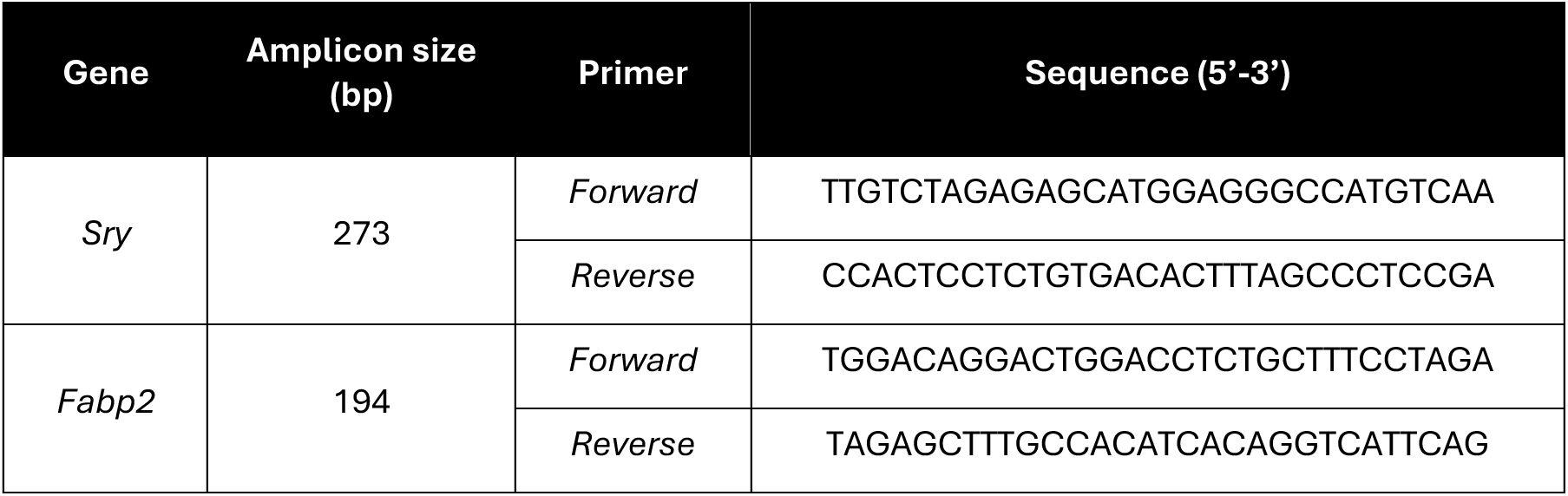
Genotyping primers for embryonic sex determination.

**Supplementary Table 4.** Immunostaining conditions and reagents.

| Experiment | Antigen Retrieval | Primary Antibody (Dilution)<br>Vendor- cat. no.,<br>RRID | Isotype Control (Dilution)<br>Vendor- cat. no.,<br>RRID | Secondary Antibody (Dilution)<br>Vendor- cat. no.,<br>RRID |
| --- | --- | --- | --- | --- |
| Spiral artery remodelling | Tris-EDTA<br>pH 9.0<br>(95°C,<br>30 min) | Mouse anti- α-SMA (1:300)<br>Abcam – ab7187<br>AB_262054<br>Rabbit anti-VIM (1:300)<br>Abcam – ab92547<br>AB_10562134<br>Rat anti- KI-67 (1:300)<br>TFS - 14-5698-82<br>AB_10854564 | Mouse IgG2a (1:300)<br>TFS - 02-6200<br>AB_2532943<br>Rabbit IgG (1:1500)<br>TFS - 02-6102,<br>AB_2532938<br>Rat IgG2a (1:600)<br>TFS - 02-9688<br>AB_2532970 | Goat anti-Mouse IgG AF488<br>TFS - A11006<br>AB_2534074<br>Goat anti-Rabbit IgG - AF555<br>TFS - A32732<br>AB_2633281<br>Goat anti-Rat IgG - AF647<br>TFS - A-21247<br>AB_141778 |
| Decidual macrophages | Tris-EDTA<br>pH 9.0<br>(95°C,<br>30 min) | Rat anti- KI-67 (1:300)<br>TFS - 14-5698-82<br>AB_10854564<br>Rabbit anti-F4/80 (1:200)<br>CST - 70076,<br>AB_2799771 | Rat IgG2a (1:300)<br>TFS - 02-9688<br>AB_2532970<br>Rabbit IgG (1:1000)<br>TFS - 02-6102,<br>AB_2532938 | Goat anti-Rat IgG - AF555<br>TFS - A48263<br>AB_2896332<br>Goat anti-Rabbit IgG – biotin (1:200)<br>VL - VECTPK6101<br>Streptavidin AF647 (1:500)<br>TFS - S32357 |

CST = Cell Signalling Technologies TFS = ThermoFisher Scientific VL = Vector Laboratories

**Supplementary Table 5.** Mouse Proteome Profiler array of E10.5 decidual tissue.

| Coordinates | Analyte | Protein Name | Relevant Aliases | Background Corrected Volume<br>(Densitometric Units) |  |  |  |
| --- | --- | --- | --- | --- | --- | --- | --- |
|  |  |  |  | CON<br>Female | HFHS<br>Female | CON<br>Male | HFHS<br>Male |
| A1/2 | - | positive control | - |  |  |  |  |
| A3/4 | AdipoQ | Adiponectin | Acrp30 | 80861.5 | 120929.5 | 120253.5 | 158433.0 |
|  |  |  |  | 78557.5 | 121430.5 | 115762.5 | 145196.0 |
| A5/6 | AREG | Amphiregulin | - | 9142.5 | 6739.5 | 7318.5 | 12768.0 |
|  |  |  |  | 8105.5 | 5765.5 | 6946.5 | 10473.0 |
| A7/8 | ANGPT1 | Angiopoetin-1 | ANG1 | 7090.5 | 9447.5 | 8956.5 | 14097.0 |
|  |  |  |  | 6306.5 | 11104.5 | 10661.5 | 16075.0 |
| A9/10 | ANGPT2 | Angiopoetin-2 | ANG2 | 389078.5 | 461713.5 | 383864.5 | 441605.0 |
|  |  |  |  | 371580.5 | 472975.5 | 386740.5 | 406699.0 |
| A11/12 | ANGPTL3 | Angiopoietin-like 3 | ANG5 | 23294.5 | 26328.5 | 38535.5 | 33415.0 |
|  |  |  |  | 13198.5 | 21265.5 | 18636.5 | 28057.0 |
| A13/14 | BAFF | B-cell Activating<br>Factor | TNFSF13B | 9276.5 | 4444.5 | 1991.5 | 9108.0 |
|  |  |  |  | 7826.5 | 7350.5 | 6284.5 | 10111.0 |
| A15/16 | CD93 | Cluster of<br>differentiation 93 | C1QR1 | 12075.5 | 35499.5 | 45084.5 | 47663.0 |
|  |  |  |  | 19719.5 | 37988.5 | 50648.5 | 36680.0 |
| A17/18 | CCL2 | chemokine C-C<br>motif ligand 2 | MCP-1 | 3520.5 | 11086.5 | 10664.5 | 12336.0 |
|  |  |  |  | 5624.5 | 11667.5 | 7534.5 | 6139.0 |
| A19/20 | CCL3/4 | chemokine C-C<br>motif ligand 3/4 | MIP-1 alpha/beta | 10848.5 | 9613.5 | 10599.5 | 9868.0 |
|  |  |  |  | 7657.5 | 9295.5 | 9656.5 | 6275.0 |
| A21/22 | CCL5 | chemokine C-C<br>motif ligand 5 | RANTES | 3006.5 | 2863.5 | 1724.5 | 505.0 |
|  |  |  |  | 3771.5 | 3208.5 | 2831.5 | 7696.0 |
| A23/24 | - | positive control | - |  |  |  |  |
| B1/2 | - | - | - |  |  |  |  |
| B3/4 | CCL6 | chemokine C-C<br>motif ligand 6 | - | 9051.5 | 13782.5 | 18814.5 | 31413.0 |
|  |  |  |  | 7132.5 | 13628.5 | 17022.5 | 27595.0 |
| B5/6 | CCL11 | chemokine C-C<br>motif ligand 11 | Eotaxin-1 | 10480.5 | 13102.5 | 13417.5 | 17390.0 |
|  |  |  |  | 7987.5 | 11878.5 | 12214.5 | 15006.0 |
| B7/8 | CCL12 | chemokine C-C<br>motif ligand 12 | MCP-5 | 9314.5 | 23571.5 | 13687.5 | 28920.0 |
|  |  |  |  | 10423.5 | 15764.5 | 17709.5 | 27995.0 |
| B9/10 | CCL17 | chemokine C-C<br>motif ligand 17 | TARC | 16814.5 | 20846.5 | 14648.5 | 33681.0 |
|  |  |  |  | 23070.5 | 19831.5 | 18485.5 | 34413.0 |
| B11/12 | CCL19 | chemokine C-C<br>motif ligand 19 | MIP-3 beta | 13169.5 | 11993.5 | 8790.5 | 22648.0 |
|  |  |  |  | 14138.5 | 11459.5 | 10395.5 | 20045.0 |
| B13/14 | CCL20 | chemokine C-C<br>motif ligand 20 | MIP-3 alpha | 1263.5 | 4079.5 | 5785.5 | 7504.0 |
|  |  |  |  | 2037.5 | 2576.5 | 7846.5 | 6710.0 |
| B15/16 | CCL21 | chemokine C-C<br>motif ligand 21 | 6CKine | 9431.5 | 8917.5 | 7843.5 | 8065.0 |
|  |  |  |  | 6518.5 | 9384.5 | 8514.5 | 11042.0 |
| B17/18 | CCL22 | chemokine C-C<br>motif ligand 22 | MDC | 34021.5 | 35233.5 | 33498.5 | 30536.0 |
|  |  |  |  | 34651.5 | 34187.5 | 37668.5 | 27456.0 |
| B19/20 | CD14 | Cluster of<br>differentiation 14 | - | 24719.5 | 31561.5 | 27631.5 | 20749.0 |
|  |  |  |  | 26291.5 | 27392.5 | 25937.5 | 23506.0 |
| B21/22 | CD40 | Cluster of<br>differentiation 40 | TNFRSF5 | 42005.5 | 36688.5 | 35150.5 | 40282.0 |
|  |  |  |  | 35141.5 | 50746.5 | 32942.5 | 37725.0 |
| B23/24 | - | - | - |  |  |  |  |
| B25/26 | - | - | - |  |  |  |  |
| C3/4 | CD160 | Cluster of differentiation 160 | - | 8633.5 | 11791.5 | 14273.5 | 14266.0 |
|  |  |  |  | 9187.5 | 9976.5 | 10303.5 | 16381.0 |
| C5/6 | RARRES2 | Retinoic acid responder protein 2 | Chemerin, TIG2 | 58434.5 | 70653.5 | 81217.5 | 101014.0 |
|  |  |  |  | 62524.5 | 78354.5 | 76747.5 | 101759.0 |
| C7/8 | CHi3L1 | Chitinase 3-like 1 | - | 7177.5 | 10606.5 | 7896.5 | 12144.0 |
|  |  |  |  | 11376.5 | 11461.5 | 9879.5 | 13151.0 |
| C9/10 | F3 | Tissue Factor | TF, CD142 | 36235.5 | 100423.5 | 37525.5 | 61022.0 |
|  |  |  |  | 40038.5 | 100062.5 | 39611.5 | 64464.0 |
| C11/12 | C5a | Complement component 5a | C5 | 6575.5 | 12869.5 | 8527.5 | 9436.0 |
|  |  |  |  | 5069.5 | 5279.5 | 5652.5 | 8717.0 |
| C13/14 | CFD | Complement factor D | Adipsin | 23405.5 | 14671.5 | 31526.5 | 20862.0 |
|  |  |  |  | 23364.5 | 15611.5 | 29987.5 | 22888.0 |
| C15/16 | CRP | C-reactive protein | - | 69923.5 | 68523.5 | 61921.5 | 78537.0 |
|  |  |  |  | 67489.5 | 69125.5 | 73022.5 | 78229.0 |
| C17/18 | CX3CL1 | chemokine C-X3-3-motif ligand 1 | Fracktalkine | 40986.5 | 36939.5 | 37494.5 | 43502.0 |
|  |  |  |  | 46360.5 | 36614.5 | 30488.5 | 44185.0 |
| C19/20 | CXCL1 | chemokine C-X-C motif ligand 1 | KC, GRO-alpha | 6306.5 | 4833.5 | 2948.5 | 736.0 |
|  |  |  |  | 4099.5 | 5994.5 | 1183.5 | 496.0 |
| C21/22 | CXCL2 | chemokine C-X-C motif ligand 2 | GRO-beta, MIP-2 alpha | 8488.5 | 11692.5 | 4742.5 | 8818.0 |
|  |  |  |  | 8725.5 | 10088.5 | 6846.5 | 9127.0 |
| C23/24 | - | - | - |  |  |  |  |
| D1/2 | CXCL9 | chemokine C-X-C motif ligand 9 | MIG | 9325.5 | 14406.5 | 10738.5 | 14413.0 |
|  |  |  |  | 5227.5 | 12796.5 | 10318.5 | 9432.0 |
| D3/4 | CXCL10 | chemokine C-X-C motif ligand 10 | IP-10 | 17972.5 | 15428.5 | 13107.5 | 17122.0 |
|  |  |  |  | 14856.5 | 18397.5 | 11103.5 | 13967.0 |
| D5/6 | CXCL11 | chemokine C-X-C motif ligand 11 | I-TAC | 4369.5 | 8659.5 | 5122.5 | 9713.0 |
|  |  |  |  | 4527.5 | 8927.5 | 7310.5 | 12211.0 |
| D7/8 | CXCL13 | chemokine C-X-C motif ligand 13 | BLC, BCA-1 | 4970.5 | 6406.5 | 7344.5 | 14522.0 |
|  |  |  |  | 3424.5 | 5550.5 | 8345.5 | 8584.0 |
| D9/10 | CXCL16 | chemokine C-X-C motif ligand 16 | - | 7646.5 | 11645.5 | 9502.5 | 15280.0 |
|  |  |  |  | 9638.5 | 15218.5 | 13576.5 | 15913.0 |
| D11/12 | CST3 | Cystatin 3 | - | 111203.5 | 117965.5 | 99910.5 | 120778.0 |
|  |  |  |  | 121786.5 | 123176.5 | 107649.5 | 115723.0 |
| D13/14 | DKK-1 | Dickkopf-related protein 1 | - | 12410.5 | 16381.5 | 16155.5 | 22934.0 |
|  |  |  |  | 20025.5 | 20521.5 | 20785.5 | 19944.0 |
| D15/16 | DPP4 | Dipeptidyl peptidase 4 | DPPIV, CD26 | 56215.5 | 80923.5 | 49498.5 | 49572.0 |
|  |  |  |  | 55170.5 | 72847.5 | 44961.5 | 48459.0 |
| D17/18 | EGF | Epidermal growth factor | - | 13186.5 | 8246.5 | 8246.5 | 11603.0 |
|  |  |  |  | 13425.5 | 11311.5 | 10840.5 | 18894.0 |
| D19/20 | Endoglin | - | ENG, CD105 | 314407.5 | 297464.5 | 308979.5 | 287168.0 |
|  |  |  |  | 299580.5 | 270339.5 | 307585.5 | 308407.0 |
| D21/22 | Endostatin | - | - | 36544.5 | 43248.5 | 40062.5 | 38362.0 |
|  |  |  |  | 37379.5 | 47988.5 | 36880.5 | 43812.0 |
| D23/24 | AHSG | Alpha-2-HS-glycoprotein | Fetuin A | 157949.5 | 158852.5 | 136684.5 | 172560.0 |
|  |  |  |  | 178181.5 | 175551.5 | 142584.5 | 185271.0 |
| E1/2 | FGF-1 | Fibroblast growth factor 1 | aFGF | 5194.5 | 7951.5 | 5026.5 | 6916.0 |
|  |  |  |  | 3199.5 | 8564.5 | 6814.5 | 11579.0 |
| E3/4 | FGF-21 | Fibroblast growth factor 21 | - | 9362.5 | 14865.5 | 8969.5 | 10915.0 |
|  |  |  |  | 9248.5 | 11081.5 | 7916.5 | 12745.0 |
| E5/6 | FLT3LG | FLT-3 Ligand | FLT3L | 4712.5 | 10709.5 | 6450.5 | 10840.0 |
|  |  |  |  | 5673.5 | 8992.5 | 11333.5 | 15273.0 |
| E7/8 | GAS6 | Growth arrest-specific 6 | - | 9851.5 | 11344.5 | 6779.5 | 14310.0 |
|  |  |  |  | 5143.5 | 14003.5 | 8994.5 | 14783.0 |
| E9/10 | CSF3 | Granulocyte colony stimulating Factor | G-CSF | 8924.5 | 3848.5 | 3448.5 | 3795.0 |
|  |  |  |  | 9067.5 | 7560.5 | 1214.5 | 8663.0 |
| E11/12 | GDF15 | Growth/differentiation factor 15 | - | 10094.5 | 12491.5 | 5023.5 | 9655.0 |
|  |  |  |  | 6258.5 | 7108.5 | 6238.5 | 13419.0 |
| E13/14 | CSF2 | Granulocyte macrophage colony stimulating factor | GM-CSF | 1207.5 | 3143.5 | 1114.5 | 4896.0 |
|  |  |  |  | 3018.5 | 4631.5 | 2888.5 | 7989.0 |
| E15/16 | HGF | Hepatocyte growth factor | - | 12811.5 | 12494.5 | 17514.5 | 17720.0 |
|  |  |  |  | 13067.5 | 13805.5 | 14915.5 | 16391.0 |
| E17/18 | ICAM-1 | Intercellular adhesion molecule 1 | CD54 | 234488.5 | 225335.5 | 235282.5 | 213990.0 |
|  |  |  |  | 236573.5 | 210987.5 | 223246.5 | 206951.0 |
| E19/20 | IFN-γ | Interferon gamma | - | 11004.5 | 15707.5 | 10406.5 | 18423.0 |
|  |  |  |  | 12667.5 | 13113.5 | 12374.5 | 16265.0 |
| E21/22 | IGFBP-1 | IGF binding protein 1 | - | 28675.5 | 19569.5 | 24307.5 | 27428.0 |
|  |  |  |  | 27874.5 | 28399.5 | 23658.5 | 25129.0 |
| E23/24 | IGFBP-2 | IGF binding protein 2 | - | 24304.5 | 29516.5 | 24127.5 | 32593.0 |
|  |  |  |  | 21198.5 | 36157.5 | 24932.5 | 35479.0 |
| F1/2 | IGFBP-3 | IGF binding protein 3 | - | 68182.5 | 74218.5 | 77254.5 | 80954.0 |
|  |  |  |  | 66662.5 | 83695.5 | 77669.5 | 80217.0 |
| F3/4 | IGFBP-5 | IGF binding protein 5 | - | 15618.5 | 20887.5 | 17944.5 | 20562.0 |
|  |  |  |  | 17062.5 | 21454.5 | 20414.5 | 24228.0 |
| F5/6 | IGFBP-6 | IGF binding protein 6 | - | 51401.5 | 45367.5 | 52861.5 | 58956.0 |
|  |  |  |  | 58648.5 | 48834.5 | 67693.5 | 57798.0 |
| F7/8 | IL-1a | Interleukin 1 alpha | - | 7935.5 | 11160.5 | 10729.5 | 12285.0 |
|  |  |  |  | 9441.5 | 10474.5 | 13376.5 | 15544.0 |
| F9/10 | IL-1b | Interleukin 1 beta | - | 6041.5 | 8399.5 | 8102.5 | 5872.0 |
|  |  |  |  | 3096.5 | 6854.5 | 5121.5 | 8069.0 |
| F11/12 | IL-1RA | Interleukin 1 receptor agonist | IL1RN | 12108.5 | 13894.5 | 5774.5 | 10962.0 |
|  |  |  |  | 7880.5 | 7892.5 | 10148.5 | 9331.0 |
| F13/14 | IL-2 | Interleukin 2 | - | -280.5 | 373.5 | -1213.5 | 1092.0 |
|  |  |  |  | 1071.5 | 1445.5 | 1425.5 | 3125.0 |
| F15/16 | IL-3 | Interleukin 3 | - | 1300.5 | 2892.5 | -517.5 | 2778.0 |
|  |  |  |  | 441.5 | 740.5 | 194.5 | 1503.0 |
| F17/18 | IL-4 | Interleukin 4 | - | 23758.5 | 20256.5 | 25086.5 | 32723.0 |
|  |  |  |  | 26298.5 | 31030.5 | 30959.5 | 33017.0 |
| F19/20 | IL-5 | Interleukin 5 | - | 8928.5 | 12610.5 | 12271.5 | 15600.0 |
|  |  |  |  | 13757.5 | 10516.5 | 8530.5 | 13981.0 |
| F21/22 | IL-6 | Interleukin 6 | - | 3957.5 | 193.5 | 1177.5 | 1222.0 |
|  |  |  |  | 2824.5 | 5387.5 | 1056.5 | 3448.0 |
| F23/24 | IL-7 | Interleukin 7 | - | 17801.5 | 22421.5 | 13109.5 | 26970.0 |
|  |  |  |  | 20026.5 | 21690.5 | 13839.5 | 24002.0 |
| G1/2 | IL-10 | Interleukin 10 | - | 8337.5 | 11307.5 | 10578.5 | 14531.0 |
|  |  |  |  | 8957.5 | 15027.5 | 11240.5 | 13581.0 |
| G3/4 | IL-11 | Interleukin 11 | - | 12742.5 | 25388.5 | 15444.5 | 21727.0 |
|  |  |  |  | 14525.5 | 19970.5 | 16682.5 | 19313.0 |
| G5/6 | IL-12 p40 | Interleukin 12 p40 subunit | IL12B | 7163.5 | 10639.5 | 12116.5 | 9268.0 |
|  |  |  |  | 9948.5 | 9123.5 | 8754.5 | 12742.0 |
| G7/8 | IL-13 | Interleukin 13 | - | 6904.5 | 14690.5 | 9407.5 | 9556.0 |
|  |  |  |  | 7767.5 | 10511.5 | 7308.5 | 14754.0 |
| G9/10 | IL-15 | Interleukin 15 | - | 12964.5 | 18508.5 | 12723.5 | 19703.0 |
|  |  |  |  | 16795.5 | 18974.5 | 17118.5 | 19027.0 |
| G11/12 | IL-17a | Interleukin 17a | - | 3730.5 | 2821.5 | 890.5 | 2997.0 |
|  |  |  |  | 1750.5 | 1039.5 | -935.5 | 2329.0 |
| G13/14 | IL-22 | Interleukin 22 | - | 4399.5 | 4517.5 | 1600.5 | 4067.0 |
|  |  |  |  | 4791.5 | 5217.5 | 709.5 | 1114.0 |
| G15/16 | IL-23 | Interleukin 23 | - | 6067.5 | 8988.5 | 9832.5 | 13728.0 |
|  |  |  |  | 6216.5 | 9435.5 | 8135.5 | 9900.0 |
| G17/18 | IL-27 p28 |  | - | 12218.5 | 17693.5 | 14764.5 | 22670.0 |
|  |  | Interleukin 27 p28 subunit |  | 9844.5 | 16497.5 | 13528.5 | 19358.0 |
| G19/20 | IL-28 A/B | Interleukin 28 A/B | IFNL2/3 | 62025.5 | 74871.5 | 62947.5 | 69741.0 |
|  |  |  |  | 66190.5 | 67899.5 | 72805.5 | 75656.0 |
| G21/22 | IL-33 | Interleukin 33 | - | 81070.5 | 97594.5 | 67802.5 | 105687.0 |
|  |  |  |  | 95059.5 | 102869.5 | 62985.5 | 108349.0 |
| G23/24 | LDL-R | LDL Receptor | - | 124106.5 | 126987.5 | 125237.5 | 143147.0 |
|  |  |  |  | 137422.5 | 141048.5 | 132270.5 | 161274.0 |
| H1/2 | Leptin | - | - | 12773.5 | 20541.5 | 10490.5 | 24376.0 |
|  |  |  |  | 12870.5 | 25548.5 | 11410.5 | 17702.0 |
| H3/4 | LIF | Leukemia inhibitory factor | - | 16376.5 | 24727.5 | 15082.5 | 26606.0 |
|  |  |  |  | 15530.5 | 21817.5 | 14938.5 | 29531.0 |
| H5/6 | LCN2 | Lipocalin 2 | NGAL | 9450.5 | 24486.5 | 12143.5 | 20717.0 |
|  |  |  |  | 11710.5 | 24018.5 | 10914.5 | 18314.0 |
| H7/8 | CXCL5 | chemokine C-X-C motif 5 | ENA78, LIX | 5274.5 | 14443.5 | 4687.5 | 17103.0 |
|  |  |  |  | 9079.5 | 10108.5 | 4888.5 | 10352.0 |
| H9/10 | CSF1 | Macrophage colony stimulating factor | M-CSF | 11711.5 | 18856.5 | 10597.5 | 13613.0 |
|  |  |  |  | 15111.5 | 18148.5 | 9740.5 | 20655.0 |
| H11/12 | MMP-2 | Matrix Metalloproteinase 2 | - | 30673.5 | 42256.5 | 34052.5 | 37028.0 |
|  |  |  |  | 35868.5 | 42011.5 | 34764.5 | 40512.0 |
| H13/14 | MMP-3 | Matrix Metalloproteinase 3 | - | -1805.5 | 2452.5 | 742.5 | 7966.0 |
|  |  |  |  | 2758.5 | 3604.5 | 1493.5 | -571.0 |
| H15/16 | MMP-9 | Matrix Metalloproteinase 9 | - | 2210.5 | 6213.5 | -759.5 | 7393.0 |
|  |  |  |  | 575.5 | 2566.5 | 6081.5 | 5988.0 |
| H17/18 | MPO | Myeloperoxidase | - | 22692.5 | 36261.5 | 27425.5 | 32153.0 |
|  |  |  |  | 28804.5 | 33813.5 | 26555.5 | 36442.0 |
| H19/20 | SPP-1 | Secreted phosphoprotein 1/ Osteopontin | OPN | 134452.5 | 143119.5 | 143015.5 | 174805.0 |
|  |  |  |  | 122269.5 | 147711.5 | 156084.5 | 162451.0 |
| H21/22 | OPG | Osteoprotegerin | TNFSF11b | 713036.5 | 778529.5 | 748670.5 | 752856.0 |
|  |  |  |  | 721851.5 | 816476.5 | 768920.5 | 759951.0 |
| H23/24 | TYMP | Thymidine Phosphorylase | PD-ECGF | 12171.5 | 20369.5 | 17438.5 | 21629.0 |
|  |  |  |  | 10132.5 | 13514.5 | 10760.5 | 22607.0 |
| I1/2 | PDFG-B | Platelet derived growth factor beta | - | 8715.5 | 17738.5 | 9281.5 | 16211.0 |
|  |  |  |  | 11119.5 | 19040.5 | 5940.5 | 20141.0 |
| I3/4 | SAP | Serum amyloid P component | APCS, PTX-2 | 135794.5 | 160795.5 | 141188.5 | 232371.0 |
|  |  |  |  | 141074.5 | 158505.5 | 145965.5 | 225528.0 |
| I5/6 | PTX-3 | Pentraxin 3 | - | 9793.5 | 20545.5 | 11453.5 | 18372.0 |
|  |  |  |  | 11668.5 | 20777.5 | 15168.5 | 24221.0 |
| I7/8 | POSTN | Periostin | OSF-2 | 2340.5 | 10144.5 | 3711.5 | 7116.0 |
|  |  |  |  | 4925.5 | 7143.5 | 3595.5 | 6659.0 |
| I9/10 | DLK-1 | Delta-like non-canonical Notch ligand 1 | - | 12090.5 | 23130.5 | 12416.5 | 19618.0 |
|  |  |  |  | 15172.5 | 22886.5 | 17507.5 | 23602.0 |
| I11/12 | PRL2C2 | Proliferin | PLF, PLF-1 | 206391.5 | 315075.5 | 188306.5 | 290910.0 |
|  |  |  |  | 199712.5 | 309067.5 | 190886.5 | 289053.0 |
| I13/14 | PCSK9 | Proprotein convertase subtilisin/kexin type 9 | - | 24574.5 | 30121.5 | 28575.5 | 33918.0 |
|  |  |  |  | 27473.5 | 35065.5 | 31680.5 | 34883.0 |
| I15/16 | RAGE | Receptor for advanced glycation end products | AGER | 9304.5 | 14770.5 | 11903.5 | 13666.0 |
|  |  |  |  | 9261.5 | 20526.5 | 11224.5 | 13378.0 |
| I17/18 | RBP4 | Retinol binding protein 4 | - | 53466.5 | 67376.5 | 53223.5 | 60161.0 |
|  |  |  |  | 51943.5 | 69736.5 | 56745.5 | 63537.0 |
| I19/20 | REG3G | Regenerating islet-derived protein 3 gamma | - | 21841.5 | 18211.5 | 24829.5 | 18144.0 |
|  |  |  |  | 13307.5 | 23467.5 | 23614.5 | 20443.0 |
| I21/22 | Resistin | - | RETN, ADSF, XCP1 | 11242.5 | 11713.5 | 10012.5 | 18292.0 |
|  |  |  |  | 7054.5 | 11537.5 | 5835.5 | 19975.0 |
| I23/24 | - | - | - |  |  |  |  |
| J1/2 | - | positive control | - |  |  |  |  |
| J3/4 | E-Selectin | - | CD62E | 7587.5 | 18133.5 | 7425.5 | 29414.0 |
|  |  |  |  | 7870.5 | 13528.5 | 10307.5 | 24017.0 |
| J5/6 | P-selectin | - | CD62P | 10376.5 | 25749.5 | 18171.5 | 32764.0 |
|  |  |  |  | 10599.5 | 35585.5 | 28526.5 | 40054.0 |
| J7/8 | PAI-1 | Plasminogen<br>activator inhibitor 1 | SERPINE1 | 344739.5 | 368942.5 | 438254.5 | 552098.0 |
|  |  |  |  | 335128.5 | 391580.5 | 402150.5 | 565750.0 |
| J9/10 | PEDF | Pigment epithelium-<br>derived growth<br>factor | SERPINF1 | 14934.5 | 20682.5 | 12791.5 | 17730.0 |
|  |  |  |  | 10792.5 | 15895.5 | 15805.5 | 18393.0 |
| J11/12 | THPO | Thrombopoetin | - | 4211.5 | 13702.5 | 7897.5 | 16156.0 |
|  |  |  |  | 8150.5 | 10260.5 | 4347.5 | 20574.0 |
| J13/14 | TIM-1 | T-cell<br>immunoglobulin and<br>mucin domain 1 | HAVCR1, KIM-1 | 51561.5 | 66557.5 | 43126.5 | 60982.0 |
|  |  |  |  | 60773.5 | 67276.5 | 50415.5 | 65231.0 |
| J15/16 | TNF | Tumor Necrosis<br>Factor | TNFa | 11080.5 | 15381.5 | 10971.5 | 17092.0 |
|  |  |  |  | 9646.5 | 16578.5 | 11921.5 | 18810.0 |
| J17/18 | VCAM-1 | Vascular cell<br>adhesion protein 1 | CD106 | 13801.5 | 16440.5 | 19851.5 | 21792.0 |
|  |  |  |  | 17443.5 | 16386.5 | 16557.5 | 19408.0 |
| J19/20 | VEGF | Vascular<br>endothelial growth<br>factor | - | 6148.5 | 21648.5 | 9132.5 | 18554.0 |
|  |  |  |  | 6353.5 | 18908.5 | 9303.5 | 15481.0 |
| J21/22 | CCN4 | WNT1-inducible<br>signaling pathway<br>protein 1 | WISP-1 | 6451.5 | 13479.5 | 7505.5 | 16059.0 |
|  |  |  |  | 5378.5 | 10845.5 | 9194.5 | 15790.0 |
| J23/24 | Negative control | Background | - |  |  |  |  |

**Supplementary Figure 1.**
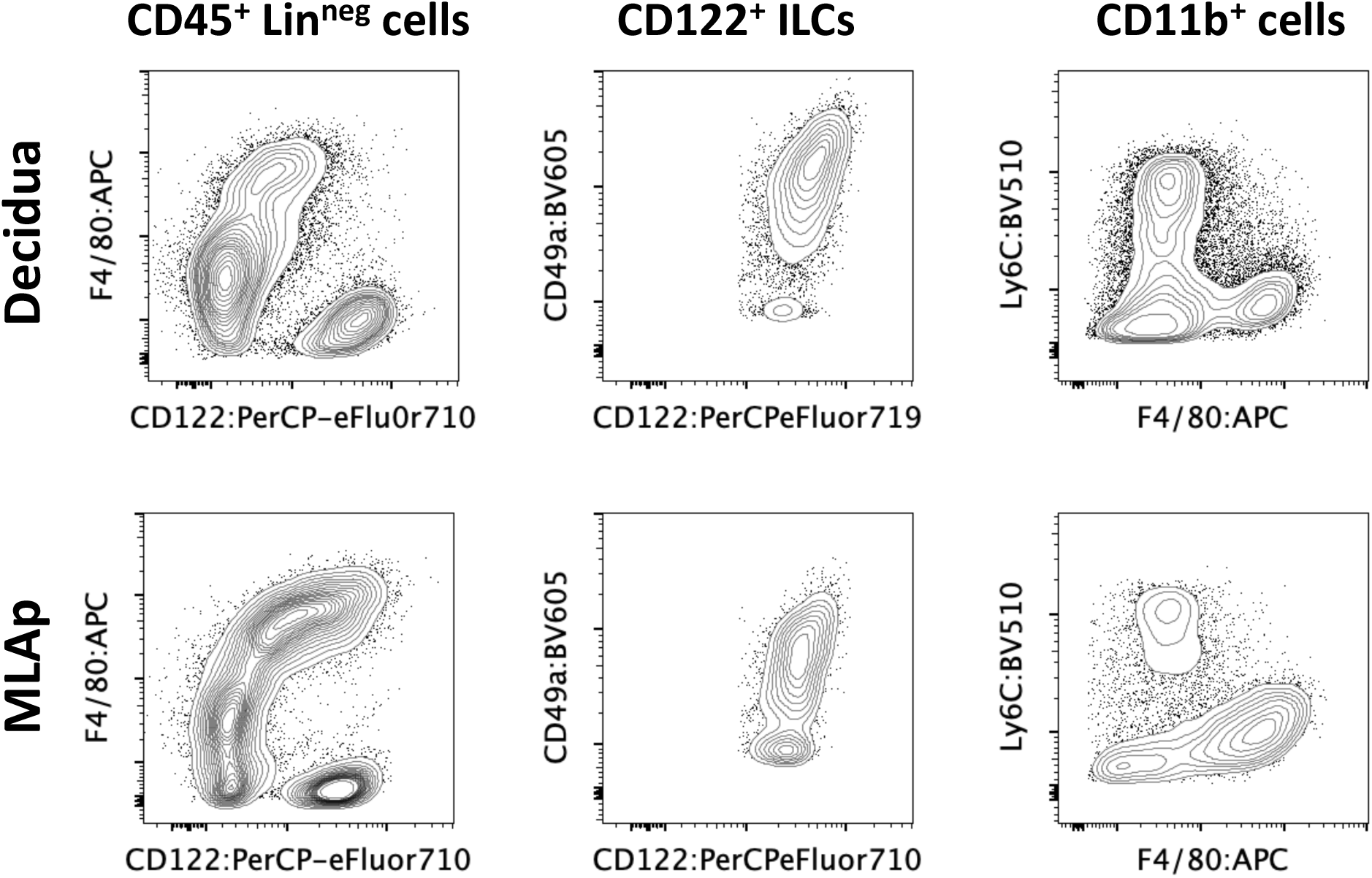
Supporting data for decidual immunophenotyping. Representative bivariate plots of flow cytometry data from a CON pregnancy at E10.5. CD45^+^ leukocytes derived from decidual (top row) or myometrium/mesometrial lymphoid aggregate (MLAp; bottom row), show distinctive profiles of staining for both innate lymphoid cell (middle) and CD11b^hi^ myeloid populations (right), indicative of proper isolation of decidual and myometrial/MLAp compartments.

**Supplementary Figure 2.**
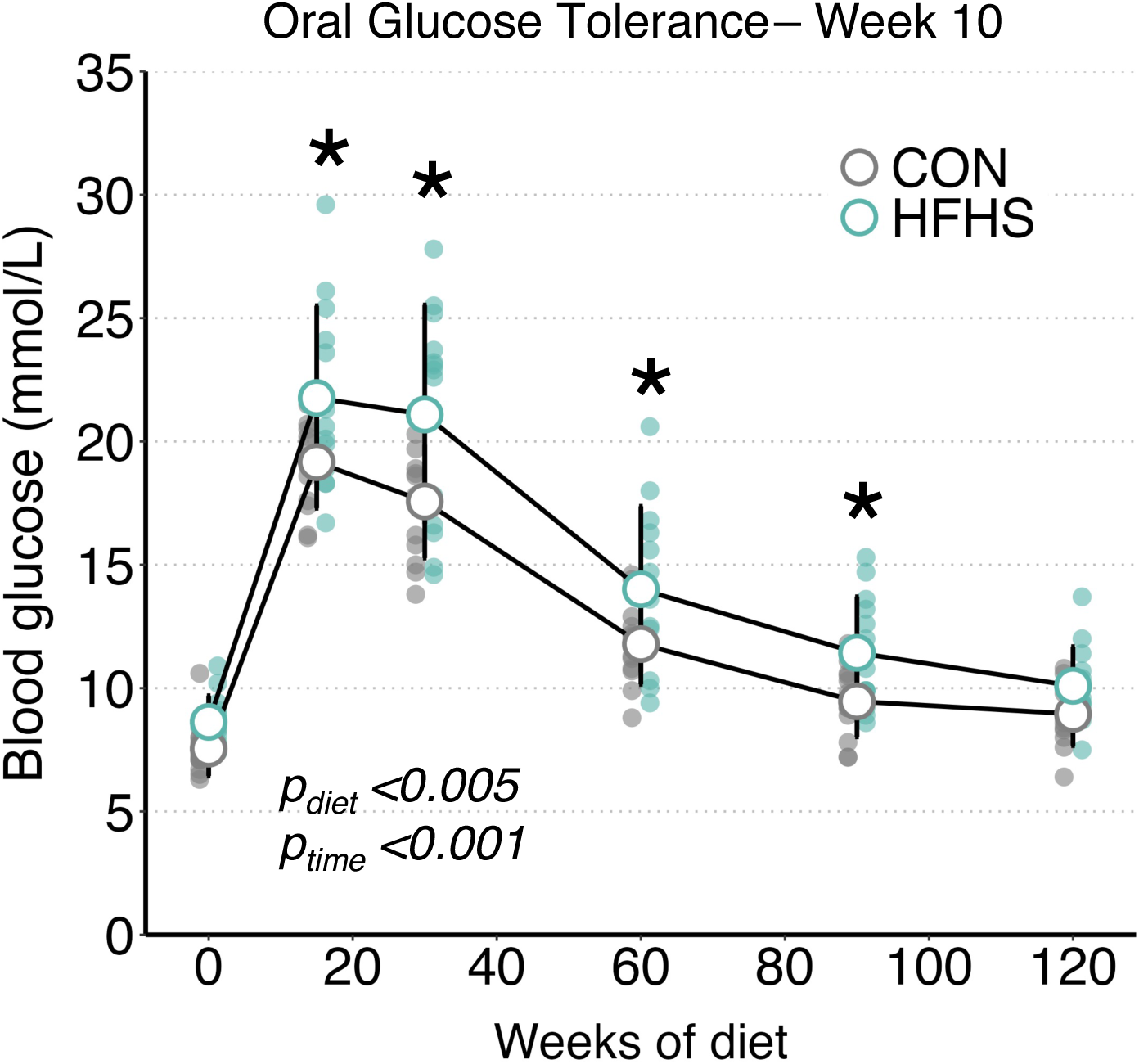
**Preconception oral glucose tolerance in terms of absolute blood glucose values**. After 10 weeks of HFHS dietary intervention, female mice were hyperglycemic during an oral glucose tolerance test but did not exhibit changes in their profile of glucose clearance (incremental area under the curve, see main Figure. Data were analysed using two-factor ANOVA on a linear mixed-effects model with diet and time as fixed effects and subject as a random effect. The main effects of diet and time are listed. Post hoc pairwise comparisons of estimated marginal means was performed by Tukey’s method at each timepoint. Data for control (CON, n =13) and high-fat, high-sucrose fed (HFHS n = 13) are presented as mean ± standard deviation (SD) underlaid with individual data points from females in each group. * Indicates p<0.05 for pairwise comparisons.

**Supplementary Figure 3.**
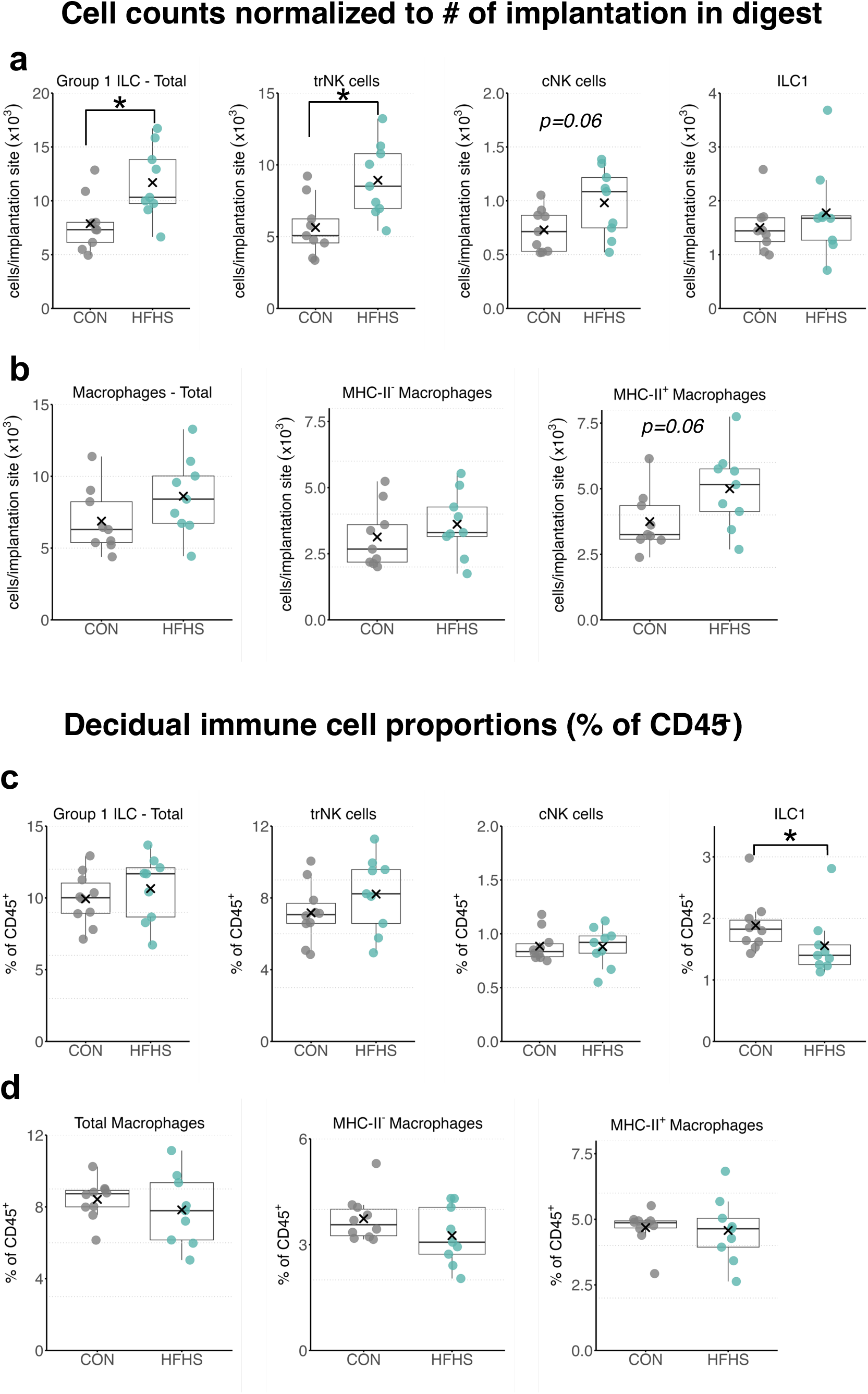
Supporting data for decidual immunophenotyping. Absolute counts of decidual **(a)** ILC and **(b)** macrophage populations in decidual cell suspensions adjusted according to the number of implantation sites used to generate samples analysed by flow cytometry (n = 3-4 implants pooled per pregnancy). The abundance of ILC **(c)** and macrophage **(d)** populations were also assessed as proportions of the total leukocyte (CD45^+^) population. Data for control (CON, n = 9) and high-fat, high-sucrose fed (HFHS n = 9) are presented as mean ± standard deviation (SD) underlaid with individual data points from females in each group in (b and h). Boxplots in (c – g, i) display IQR (box), median (line), with whiskers indicating min/max values for each group within 1.5 SD of the mean (denoted by ‘X). Pairwise comparisons were made using linear regression with maternal diet as the independent variable. * Indicates p<0.05 for pairwise comparisons.

**Supplementary Figure 4.**
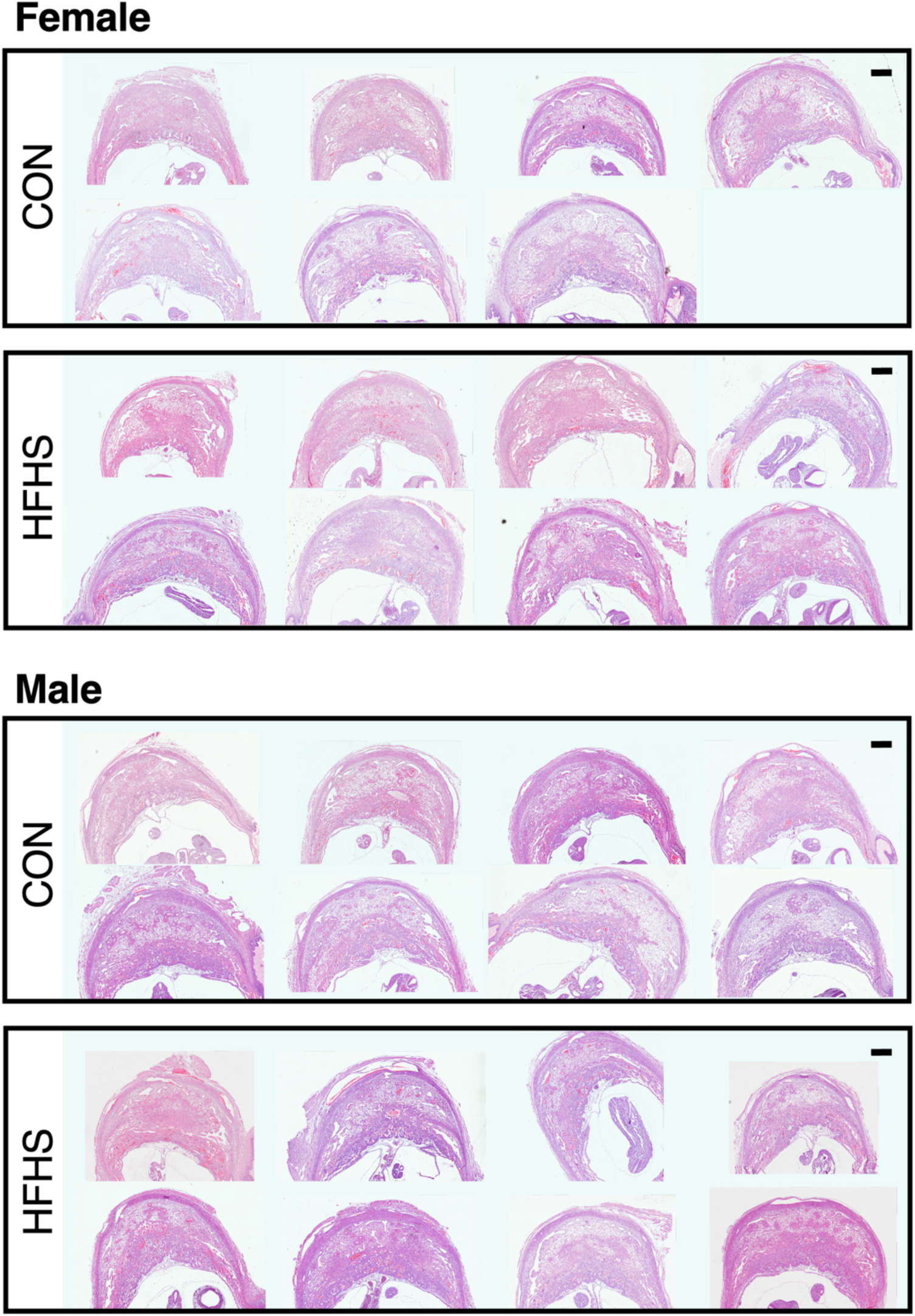
Implantation site morphology is macroscopically similar between CON and HFHS pregnancies. Mid-sagittal tissue sections of H&E-stained implantation sites from CON (n=7 – 8/sex) and HFHS (n = 8/sex) pregnancies. Composite images were generated from fields imaged at 20× magnification. Scale bar = 500 μm.

